# The characterization of a humanized rabbit anti-TNFα monoclonal antibody

**DOI:** 10.1101/2019.12.18.881029

**Authors:** Chao Wang, Xianpu Ni, Ying Liu, Zheng Jin, Huanzhang Xia

## Abstract

Tumor necrosis factor alpha (TNFα) is now regarding as a key role in the pathogenesis of immune-mediated disease such as Rheumatoid arthritis (RA), Crohn's Disease, Psoriatic arthritis and Plaque Psoriasis. HERE we have successfully developed an anti-hTNFα monoclonal rabbit antibody(HZ3M) with high binding and neutralizing activity based on RabMAbs platform. Rabbit hybridomas, immunized subcutanrously with 0.4 mg human TNFα, were generated from the rabbit splenocytes and a total of 142 hybridoma clones with specific binding to human TNFa were obtained. The anti-TNFa RabMAbs showed better neutralizing activity and higher antigen binding affinity compared to Humira and Remicade, the elimination phase half-life 58.2h respectively. In vivo efficay studies, normal mice or human TNF-alpha transgenic mice were injected with 1.0 mg/kg Humira (positive control), HZ3M at 0.33?1.0 or 3.0 mg/kg, or solvent (negative control), showed that HZ3M is able to bind and neutralize hTNFα in transgenic and normal mice as well as normal rabbits.Clearly dose-dependent response can be determined. Compared to marketed anti-TNFa drug Humira, the efficacy of HZ3M is seems to show significant longer holding time.Our observations indicate that the HZ3M derived from RabMAb preclinical safety study, and might have a therapeutic role in RA treatment.

## INTRODUCTION

Tumor necrosis factor alpha (TNFα) is now regarding as a key role in the pathogenesis of immune-mediated disease such as Rheumatoid arthritis (RA), Crohn’s Disease, Psoriatic arthritis and Plaque Psoriasis. RA is a chronic disease in which inflammation of the synovial tissue results in articular cartilage and bone destruction. The concentration of TNFα in the sites of inflammation has been found arise abnormal and drive RA pathology by direct inflammation and TNFα regulated cytokines and chemokines. The concept that the removal of TNFα in the inflammation sites can prevent the further development of RA has been confirmed by the successes of several biological TNFα antagonists: fully human or chimeric IgG_1_ monoclonal antibodies adalimumab,infliximab and golimumab. By binding to TNFα and preventing the interaction between TNFα and TNFα receptor, this kind of MAbs can diminish the concetration of diminish and neutralize the biological function as well. The therapeutic efficacies of this kind TNFα antagonists are so promising that most of them have been predicted as the blockbusters in the next 10 years. However, the applications of such anti-TNFα monoclonal antibodies have been connected with some several but rare adverse reactions such as TB infection and malignancy. The biggest one is still the immunogenicity concern especially the arise of anti-drug antibody which dramatically decrease efficacy of drug. Unlike those cases of PRCA happened in the patiences using EPO, most patients who fail to respond, have lost response or are intolerant of one TNFα antagonist respond well when switched to another TNF antagonist^1^.

The development of new anti-TNFα monoclonal antibody is not only to fulfil the increased demanding of biological TNFα antagonists, but also to provide substitutable option to those intolerant/unresponse with one certain producte. Here we use RabMAbs technology to build up a new humanized anti-human TNFα monoclonal antibody, and characterizations have been made based on in vivo pharmacokinetics and efficacies study.

## MATERIALS AND METHODS

### Generation of Anti-human TNFa RabMAbs

New Zealand rabbits were immunized subcutaneously with 0.4 mg human TNFa (Shanghai Primegene Bi-Tech) in complete adjuvant (Sigma Chemical, St. Louis, MO). After the initial immunization, animals were boosted 5 times in a 3 week-interval with 0.2 mg human TNFa in incomplete adjuvant (Sigma Chemical, St. Louis, MO). Final boost was given intravenously with 0.4 mg the protein in PBS four days before splenectomy. Hybridoma fusion was performed according to established protocol with minor modifications^2^. Briefly, splenocytes were harvested from the immunized rabbit and fused with rabbit plasmacytoma 240E-W^3^, a subcloned line of 240E^2^ using PEG4000 (Sigma Chemical, St. Louis, MO). The hybrid cells, hybridomas, were selected by HAT (hypoxanthine, aminopterin, and thymidine). At end of the selection, the medium were changed twice. Five days later, hybridoma supernatants from fusion plates were screened by direct antigen (hTNFa) binding ELISA. The positive clones were then transferred to 24-well plates and confirmed by the binding ELISA. To identify hybridoma clones producing anti-human TNFa RabMabs with neutralizing activity, supernatants from the confirmed ELISA positive clones were collected and tested in L929 TNFa mediated cell toxicity and NFkB nuclear translocation assays. The clones showing TNFa neutralizing activity in these two assays were subcloned and supernatants from subclones of the clones were screened again for the antigen binding and neutralizing of TNFa. The subclones that were positive in the binding and neutralizing assays were expanded and frozen for future use.

### Cloning and Expression of Recombinant Anti-human TNFa RabMAbs

Total RNA from Hybridomas (10^4^-10^5^ cells) producing desired RabMAb was isolated by using Qiagen RNeasy kits and annealed to oligo dT before use for reverse transcription (RT) reaction. RT reaction was carried out in 15 ul reaction mixture at 40 °C for 60 min after addition of ImProm II (Promega). DNA fragments for L chain and the variable region of H chain (VH) of rabbit IgG were amplified by PCR in 25 ul reaction mixtures containing 2ul of RT products, 1.25 units of Taq Plus Precision DNA polymerase (Stratagene), 0.2 mM dNTP and 0.3 uM of either L chain or VH primers. The L chain fragment was cloned into pCDNA 3.1 vector (Invitrogen) at BamH I and Xho I sites or pCEP4 vector (Invitrogen) at Hind III (or Kpn I) and Xho I sites, and the VH was fused in-frame to the constant region of H chain at BspE1 site and cloned into another pcDNA 3.1 or pCEP4 vector at BamH I and Pme I sites. Recombinant RabMAbs were expressed in HEK 293 or CHO cells by cotransfection of the L and H chain plasmids. For humanized anti-human TNFa RabMAb cloning and expression, the humanized VH was fused in-frame to the constant region of human IgG1 H chain at Nhe I site and clones into pTT5 vector^4^ at Hind III and Not I sites, and the humanized VK to the constant region of human IgG kappa chain at BsiWI site and into another pTT5 vector at Hind III and Not I sites. The L and H chain plasmids were co-transfected into 293-6E cells^4^ and supernatants were harvested 5 days after transfection. The recombinant RabMAbs in the supernatants were purified through protein A column, dialyzed in PBS buffer and quantified at OD280 nm.

### Antigen Binding ELISA

For the direct antigen binding ELISA assay, recombinant human TNFa (Shanghai Primegene Bi-Tech) was coated onto microtiter plates (Greiner Bio-One) in 0.05M carbonate-bicarbonate buffer overnight at 4°C, and non-specific binding was blocked with 1% BSA/TBST (Tris-Buffer Saline Tween-20) for 1 hour at room temperature. Rabbit anti-human TNFa hybridoma supernantants or recombinant antibodies were added to the plates for binding to the antigen. Alkaline phosphatase-conjugated goat anti-rabbit (Jackson ImmunoResearch) and goat anti-human IgG (H+L) antibody (Pierce/Thermo Scientific) were added at a dilution of 1:5000 to detect bound rabbit and human antibodies, respectively. The ELISA plates were then processed for pNPP substrate development and the optical density was measured at 405 nm with an ELISA plate reader (Labsystems Multiskan Ascent Photometric plate reader).

### L929 TNFa Induced Cell Cytotoxicity Assay

L929 cells (ATCC, 3.5 x 10^4^ /well) were seeded into 96-well assay plates in 100 ul of Dulbeco’s modified Eagle’s medium (Cellgro) supplemented with 2% fetal bovine serum (Hyclone, heat inactive) and incubated overnight at 37 C, 5% CO2 in a humidified incubator. Recombinant human TNFa (R&D, final assay concentration, 1ng/ml) was preincubated with or without anti-TNFa hybridoma supernatant, parental or humanized recomboinant RabMAb or control antibody at 37 C for 2 hrs, and then 50 ul of the preincubated mixture was transferred to the assay plates after addition of 50 ul of actinomycin D at a final concentration of 1ug/ml (Sigma). Incubation of the assay plate was continued for 18 h followed by addition of 20 μl of a MTT Reagent (R&D). After another 4 hrs incubation, 100 μl of Detergent Reagent (R&D) was added. Next day, the absorbance of each well was determined with Wallac 1420 Victor plate reader at 570 nm. Survival was calculated as the percentage of the staining value of untreated cultures. Percent cytotoxicity is the difference between control (100%) and percent survival.

### NFkB Nuclear Translocation Assay

Hela cells (ATCC, 1000 /well) were seeded into 96-well assay plates in 100 ul of Dulbeco’s modified Eagle’s medium (Cellgro) supplemented with 10 % fetal bovine serum (Hyclone, heat inactive) and incubated overnight at 37 C, 5% CO2 in a humidified incubator. To screen for RabMAbs that can neutralize TNFa activity in induction of NFkB nuclear translocation, the cells were untreated, treated with 1 ng/ml recombinant human TNFa (R&D) or 1 ng/ recombinant human TNFa that was preincubated for 20 min with a positive control antibody (Upstate) or hybridoma supernatant. After 20-30 min treatment, the cells were washed twice in TBST (Tris-Buffer Saline Tween-20) and fixed for 20 mins in 2% paraformaldehyde at room temperature. To stain, the cells were permeabilized for 5 mins with 0.1% Triton-100 /TBS (Tris-Buffer Saline) at room temperature, blocked for 1 hr in 1% BSA in TBS and then incubated for 1 hr with mouse anti-NF-KB p65 antibody (BD biosciences) at 1:1000 dilution in 1% BSA. After three washes in TBST, anti-mouse-HRP antibody (ImmunoVision Technologies) was added and the cells were incubated with the secondary antibody for 2 hrs. The assay plate was then processed for DAB (ImmunoVision Technology) substrate development and cell stain was observed under a microscope.

### Humanization of TNFa RabMAb

A phylogenetic tree was built by alignment of VH and VL amino acid sequences of selected RabMAbs. Sequence alignment and phylogenetic analysis were performed using the ClustalX software. The parental RabMAb and the most closely related human germline sequence were then aligned. RabMAb residues were changed to match the human germline sequence except for the residues which are known to be likely critical in antigen binding. By MLG analysis^5,6^, some of the critical residues subjected to change during the *in vivo* maturation process were identified and humanized. DNA encoding humanized VK and VH were synthesized by MCLab (South San Francisco, CA, USA). The human IgG signal peptide and a Kozak sequence were engineered at the 5’ends of the VK and VH sequences. (For the cloning and expression of humanized anti-TNFa RabMAb, see **Cloning and Expression of Recombinant Anti-human TNFa RabMAbs)**

### Thermal Stability Studies

Purified recombinant antibodies in phosphate buffer saline, pH 7.4, were subjected to thermal denaturation over the temperature range of 37- 95 °C in 1 cm path length, reduced-volume, quartz cells (800 ul). The thermal denaturation profiles, absorbance at 280 nm as a function of temperature, were acquired on a Cary 100 spectrophotometer system with accessories for temperature control. The rate of temperature increase was 1 °C/min.

### Analysis of Glycosylation on Anti-TNFa RabMAbs

Purified recombinant antibodies in glycoprotein denaturing buffer (0.5% SDS and 1% β-mercaptoethanol) were heated at 100 °C for 10 min, and then sodium phosphate, pH 7.5 and NP-40 were added to a final concentration of 50 mM and 10%, followed by addition of PNase F (New England Bioland). Deglycosylation reaction was performed at 37 C for 10 min. The glycosylated and deglycosylated proteins were separated on a 4-20% Novex Tris-Glycine gel (Invitrogen) by reducing, denatured gel electrophoresis and visualized by Gelcode (Pierce) blue stain.

### Affinity Measurement of Anti-TNFa RabMAbs by competition ELISA

The optimal coated antigen and antibody concentrations used in competition ELISA were pre-determined by direct antigen binding ELISA (see **Antigen Binding ELISA** for the detailed method). Five different hTNFa concentrations ranging from 0.02 to 0.50 ug/ml were tested for coating in 96-well plates. For each of the antigen coating concentrations, recombinant antibodies in transient expression supernatants were applied to binding to the antigen without dilution or with 1: 3, 1:9 or 1:27 dilution. The hTNFa coating concentration giving the greatest difference between maximum and minimum antibody bindings was selected for determination of the optimal antibody concentration. A serial of recombinant antibody dilutions in a 96-well plate (1:3, from row A to G; no antibody addition to row H) were tested for binding to the coated antigen. Antibody bindings vs antibody concentrations were plotted. The optimal antibody concentration was the median effective concentration, EC50, at which the antibody binding reached half maximum one.

In competition ELISA, hTNFa (15 nM) with a serial of dilutions (1:3) from row A to G in a BSA-blocked 96 well plate was incubated with an anti-TNFa antibody diluted to the optimal concentration for 30 min, and then the pre-incubated antigen/antibody mixes were transferred to a separate 96-well plate coated with hTNFa at the optimal concentration overnight and blocked with BSA. After incubation for 60 min, the plate was washed thoroughly with TBST and processed as a direct antigen binding ELISA plate. The dissociation constant (Kd) was the median inhibitory concentration, IC50, at which hTNFa in solution achieved inhibition of the antibody binding to the coated hTNFa by 50% of the maximum.

### Affinity Measurement of Anti-TNFa RabMAbs by BIAcore

Affinities measured by BIAcore were performed on BIAcore 3000 (BIAcore, Inc., Uppsala, Sweden). Antibodies were immobilized on CM5-Sepharose chips coated by standard amine coupling chemistry to a density of about 600 response units (RU) Measurements were carried out at 25°C by injecting 250 μl of recombinant human TNFa (R&D Systems) at concentrations from 0.2 to 50 nM with a flow rate of 35μl/min. The dissociation duration for bound analytes was 15 minutes in HBS-P running buffer (10 mM HEPES, pH 7.4, 150 mm NaCl and 0.005% surfactant P20, filtered and degassed). Data were analyzed with the BIAevaluation software (version 3.2RC1) by applying a 1:1 binding model. The obtained sensorgrams were fitted globally over the whole range of injected concentrations and simultaneously over the association and dissociation phases. Equilibrium dissociation constants were then calculated from the rate constants (*K_D_* = *k_off_/k_on_*).

### Pharmacokinetics study

The HZ3M (also named as HZD-RabMAb above) and anti-human TNFα RabMAb was produced as described above. Humira control was purchased from Abbott Laboratories, lot number 41366SP40. Eight weeks old male C57BL/6 mice, weight 20 ± 2g, were purchased from Vital River Laboratories. The animals were given free diet during the study.

The HZ3M, anti-TNFα RabMAb, or Humira was given to mice i.p. at 10 mg/kg. The orbital venous blood was taken from the treated animals at 5mim 30min, 1h 4h, 8h, 24h, 96h, 8d, 11d, 15d, 22d, 29d, 36d, 43d after the injection, five animals per time point. The blood was centrifuged and the serum was collected and stored at −70°C.

The serum drug concentration was measured by ELISA (enzyme-linked immunosorbent assay)^7^. The 96-well microtiter plates were coated with human recombinant TNFa (Beijing Biosea Biotechnology Co.,LTD) as the capture reagent. After the animal blood samples or Humira were incubated with the coated plate, goat-anti-human IgG-HRP (from Zhongshan Goldenbridge Biotechnology Co.,LTD) was used as the detecting reagent.

The pharmacokinetics analysis was carried by using the single-compartment open model with first order absorption to calculate elimination phase half-life^8,9^

### Efficacy Study

Testing samples: Humanized rabbit antiTNFa monoclonal antibody HZ3-M was produced and purified as described above. Humira was purchased from Abbott Laboratories, lot number 65204LJ40.

#### Mice RA study

Animals: The normal mice, C57BL/6, 6-7 weeks of age, male, were purchased from the Vital River, Corp. The human TNFa transgenic mice (Model 1006), age 5-9 weeks, male, were purchased from Taconic Corp.

Method: The animals were divided into six groups as shown in table 1. The HZ3M (resting agent), Humira (positive control), or solvent (negative control) were injected i.p. at 0.1ml/10g body weight volume.

**Table 1.** The animal grouping for the RA efficacy study

|  | Group | Treated Agents | Animals | Animal No. |
| --- | --- | --- | --- | --- |
| 1 | Normal animal control | Solvent | Normal C57BL/6 mice | ♂ 10 |
| 2 | Negative control | Solvent | Taconic 1006 mice | ♂ 10 |
| 3 | Positive control | 1mg/kg Humira | Taconic 1006 mice | ♂ 10 |
| 4 | HZ3M, low dose | 0.33mg/kg HZ3M | Taconic 1006 mice | ♂ 10 |
| 5 | HZ3M, medium-dose | 1mg/kg HZ3M | Taconic 1006 mice | ♂ 10 |
| 6 | HZ3M, high dose | 3mg/kg HZ3M | Taconic 1006 mice | ♂ 10 |

The HZ3M (testing agent), Humira (positive control), or solvent (negative control) were given to the animals by i.p. injection, once every three days for six weeks, at 0.1ml/10 g body weight volume.

The RA symptoms were analyzed and scored once a week, based on the limbs swollenness, toes curvature, toes separation, toe joints and ankle joints mal-shape^10^.

| Swollenness |  |  | Redness |  | Toe curvature |  |  |  |
| --- | --- | --- | --- | --- | --- | --- | --- | --- |
| 0=no | 0.5=mild | 1=sever | 0=no | 1=yes | 0=no | 0.3=mild | 0.7=medium | 1=severe |
| Toe separation |  |  |  |  |  | Toe joint mal-shape |  | Ankle joint mal-shape |
| 0=full | 0.5=mild | 0.4=2 toes clustered | 0.6=3 toes clustered | 0.8=4 toes clustered | 1=5 toes clustered | 0.2 for every mal-shaped toe joint |  | 0.5 for every mal-shaped ankle joint |

Mice muscle strength study: The mice were subject to the muscle strength test once a week on the Rota-Rod Instrument during the treatment. The holding time was recorded, above 60 seconds were counted as 60 second.

Joint histology analysis: The mice hind leg ankle joints, collected one week after the treatment was completed, were fixed in 10% formalin over night, decalcified in 30% formic acid, paraffin embedded, sliced, and H.E. stained. The histological sections were evaluated under microscope based on the pathology and the scored as below:

| Score | 0 | 1 | 2 | 3 | 4 |
| --- | --- | --- | --- | --- | --- |
| Symptom | no pathological change | synovial hyperplasia and leukocyte infiltration | pannus and tissue fibrosis, erosion of local cartilage lesion | articular cartilage and bone erosion | extensive articular cartilage and bone erosion |

#### Mice survival study

Animals: The normal mice, C57BL/6, female, body weight 18-22g, were purchased from the Vital River, Corp.

Treatment: The animals were divided to four groups (normal, negative control, positive control, and HZ3-M) and subjected to various treatments as shown in table 2.

**Table 2.**
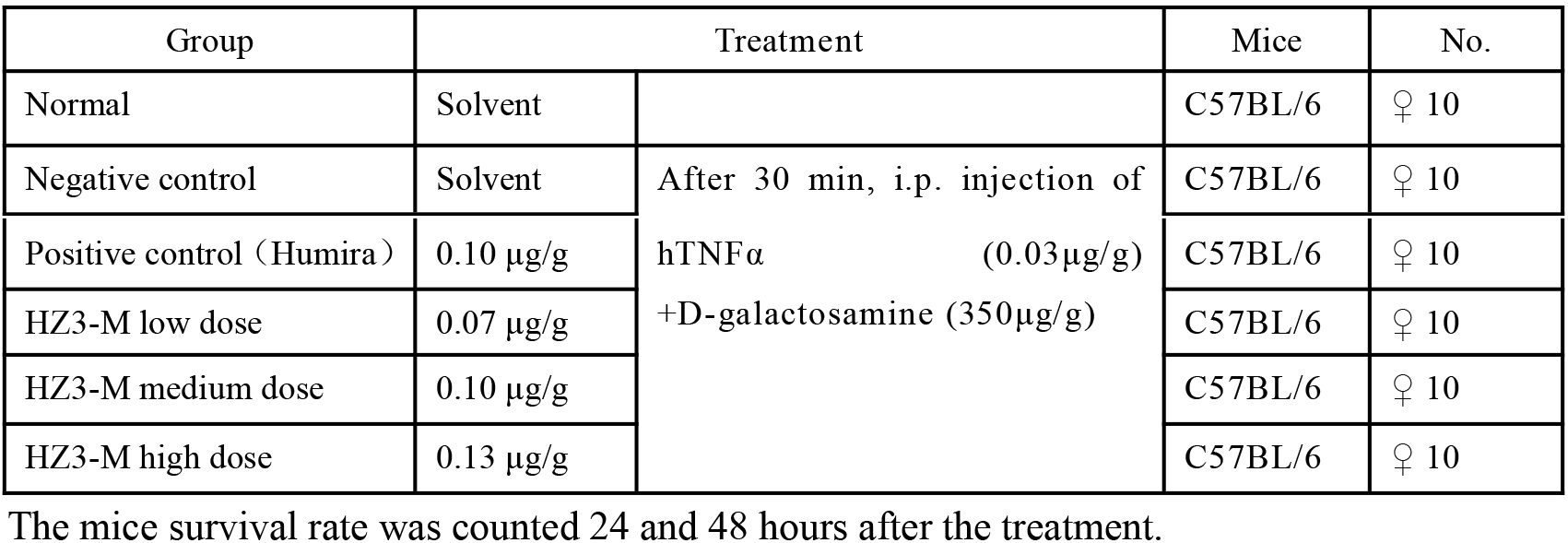
The animal grouping and treatments for the survival study

#### Fever study

Animal: Closed colony of New Zealand rabbits, 50% male and 50% female, were purchased from Shenyang Pharmaceutical University Experimental Animal Center.

Treatment: The animals were divided to four groups (normal, negative control, positive control, and HZ3-M) and subjected to various treatments via ear vein injection as shown in table 3. The animal Rectal temperature was measured every 30 min within 4 hours after the treatment.

**Table 3.** The animal grouping and treatments for the fever study

| Group | Treatment | Animal | No. |
| --- | --- | --- | --- |
| Normal | Solvent | Normal rabbit | ♂♀ 6 |
| Negative control | Solvent+TNFα (1µg/kg) +D-Gln (2µg/kg) | Normal rabbit | ♂♀ 6 |
| Positive control (Humira) | 50µg/kg+TNFα (1µg/kg) +D-Gln (2µg/kg) | Normal rabbit | ♂♀ 6 |
| HZ3-M, low dose | 25µg/kg+TNFα (1µg/kg) +D-Gln (2µg/kg) | Normal rabbit | ♂♀ 6 |
| HZ3-M, medium dose | 50µg/kg+TNFα (1µg/kg) +D-Gln (2µg/kg) | Normal rabbit | ♂♀ 6 |
| HZ3-M, high dose | 100µg/kg+TNFα (1µg/kg) +D-Gln (2µg/kg) | Normal rabbit | ♂♀ 6 |

## RESULTS

### Generation of Anti-human TNFa RabMAbs with Neutralizing Activity

Three rabbits were immunized with full length human TNFa expressed in E. coli. Antisera from the rabbits were tested for their activity to neutralize TNFa induced cytotoxicity in L929 cells. Antisera from all the rabbits showed the neutralizing activity (Fig. 1). Rabbit hybridomas were generated from the rabbit splenocytes and a total of 142 hybridoma clones with specific binding to human TNFa were obtained. Forty-four RAbMAbs that neutralize TNFa induced cytotoxicity in L929 cells were identified from screening of the 142 clones in the cell-based assay (Fig 2a). The neutralizing activity was confirmed by their capability to block NFkB nuclear translocation (Fig 2b). To rank the neutralizing antibodies, cDNAs for the neutralizing antibodies were clones and expressed and dose-responsive neutralization of TNFa-induced cytotoxicity assay was performed with the recombinant antibodies. IC50 was calculated and summarized in Table 4. Antigen binding affinity for some of these antibodies was also measured (Table 4). Currently marketed anti-TNFa drugs, Humira and Remicade, were used as controls. Some of anti-TNFa RabMAbs showed better neutralizing activity and higher antigen binding affinity compared to Humira and Remicade.

**Fig.1.**
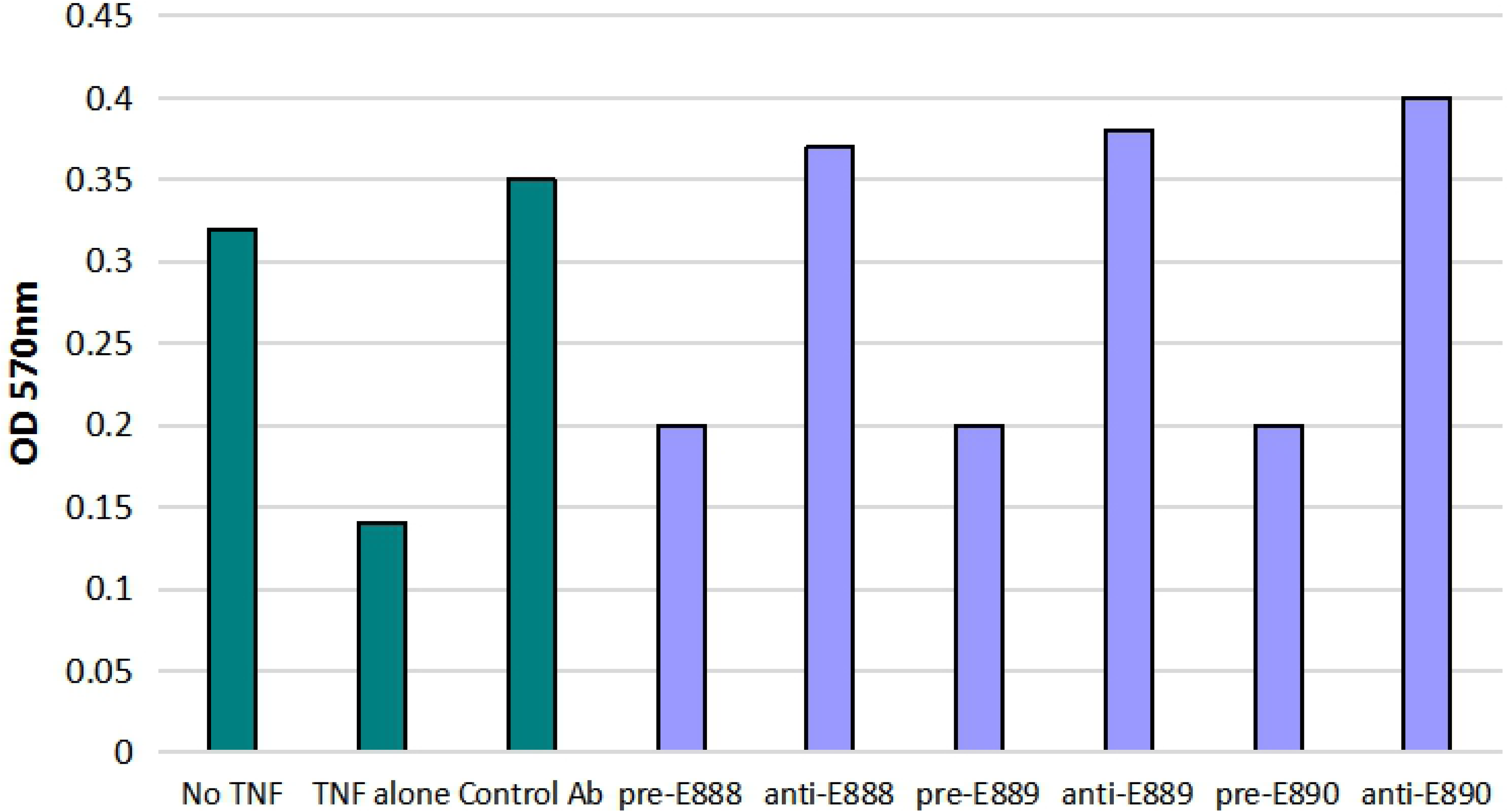
Neutralizing activity of recombinant human TNFa immunized rabbit sera. Rabbit sera from three TNFa immunized rabbits (anti-E888, −889 and −890) and from corresponding rabbits prior to immunization (pre-E888, −E889 and −890) were tested at 1:12, 800 dilution in TNFa-induced L929 cell cytotoxicity assay. After incubation of the cells in 96-well plates with the sera, MTT assay was performed and absorbance at 570 nM was measured. The cells without (No TNF) or with (TNF alone) TNFa treatment or with TNFa treatment plus control antibody (Cntrl Ab) were run in parallel.

**Fig. 2a.**
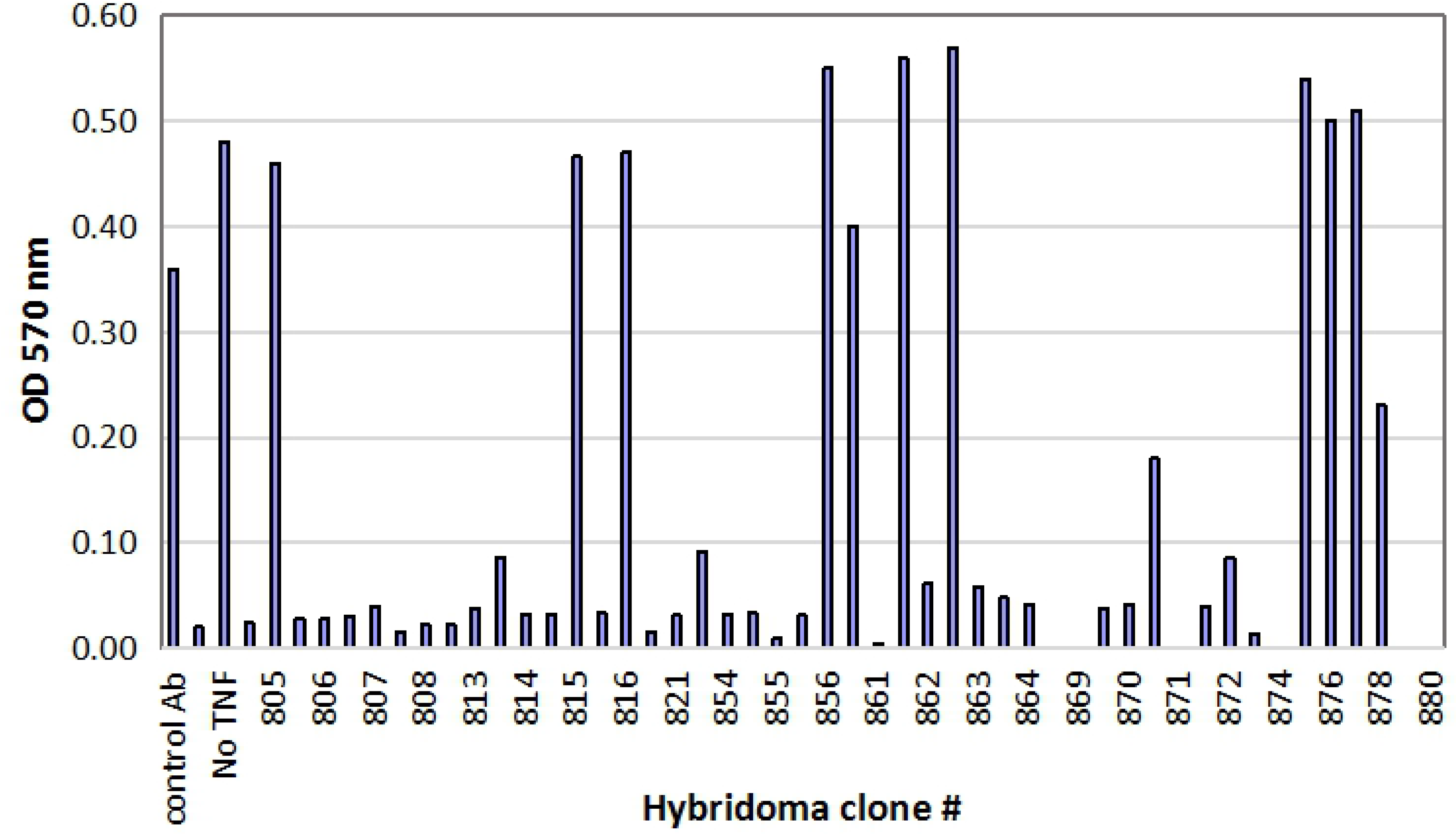
Representative data from screening of hybridoma supernatants by TNFa induced cytotoxicity assay. L929 cells in 96-well plates were treated with TNFa and anti-TNFa RabMAbs in hybridoma supernatants from different clones, and then MTT assay was performed and absorbance at 570 nm was measured. The cells without (No TNF) or with (TNF alone) TNFa treatment or with TNFa treatment plus control antibody (Cntrl Ab) were run in parallel.

**Fig. 2b.**
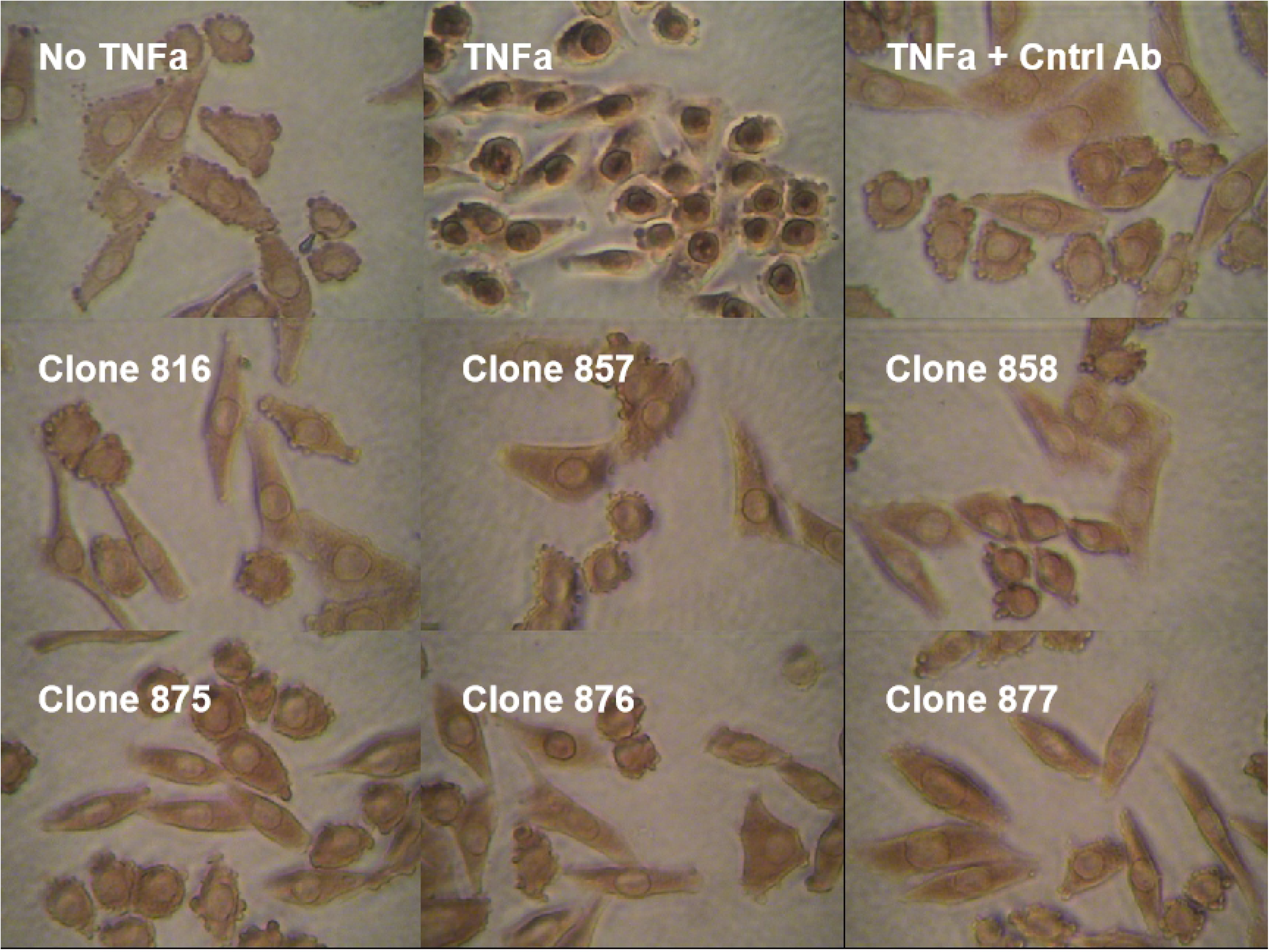
Representative data from screening of hybridoma supernatants by NFkB nuclear translocation assay. Hela cells were treated with TNFa and anti-TNFa RabMAbs in hybridoma supernatants from different clones and then fixed with Paraformaldehyde. NFkB in the cells was probed with mouse anti-NFkB antibodya and its location was detected by anti-mouse HRP-polymer and DAB stain. The cells without (No TNF) or with (TNF alone) TNFa treatment or with TNFa treatment plus control antibody (Cntrl Ab) were run in parallel.

**Table 4.** Neutralizing activity and binding affinity of anti-TNFa RabMAbs

| Clone # | Neutralizing Activity IC50 (nM) | Binding Affinity Kd (nM) | Clone # | Neutralizing Activity IC50 (nM) | Binding Affinity Kd (nM) | Clone # | Neutralizing Activity IC50 (nM) | Binding Affinity Kd (nM) |
| --- | --- | --- | --- | --- | --- | --- | --- | --- |
| 878 | 763 | n.d. | 710 | 173 | n.d. | 311 | 1628 | n.d. |
| 877 | 123 | 0.15 | 703 | 97 | n.d. | 204 | 100 | 1.00 |
| 876 | 47 | 0.63 | 518 | 300 | n.d. | 115 | 210 | 0.65 |
| 875 | 80 | n.d. | 502 | 727 | n.d. | 101 | 137 | 0.15 |
| 867 | 1029 | n.d. | 473 | 130 | 3.00 | 92 | 287 | 0.20 |
| 859 | 170 | n.d. | 442 | 317 | n.d. | 89 | 577 | 0.15 |
| 858 | 147 | 0.16 | 438 | 800 | n.d. | 63 | 87 | n.d. |
| 857 | 553 | 0.24 | 433 | 667 | n.d. | 52 | 37 | 2.00 |
| 830 | 128 | 0.53 | 431 | 463 | n.d. | 48 | 1270 | n.d. |
| 825 | 250 | n.d. | 427 | 858 | n.d. | 45 | 453 | 2.00 |
| 818 | 228 | 0.21 | 410 | 1007 | n.d. | 14 | 165 | n.d. |
| 816 | 137 | 0.50 | 406 | 1390 | n.d. | 12 | 313 | 0.25 |
| 815 | 153 | 0.19 | 402 | 344 | n.d. | 1 | 113 | 4.00 |
| 810 | 217 | 0.20 | 325 | 160 | n.d. | Humira | 127 | 0.30 |
| 715 | 40 | 0.20 | 317 | 1507 | n.d. | Remicade | 169 | 0.50 |

### Sequence and Structure Analysis of Anti-TNFa Neutralizing RabMAbs

Phylogenetic trees based on variable region amino acids sequences of light (VK) and heavy (VH) chains of the neutralizing RabMAb are shown in Figure 3a. There are seven lineage-related groups identified. In the same lineage-related group, all the VHs are in the same cluster, so as all the VKs. Sequence alignments of the lineage related groups are shown in Figure 3b (VH) and 3c (VK). RabMAbs in a lineage-related group have the same length of CDRs and their sequences in the heavy chain CDR3 are very similar as well. For some lineage groups, the CDR2 sequence is more diverse than the CDR3 in the heavy chains (Fig. 2b Group A, B and G). In addition, some significant sequence variations were observed in the CDR1, CDR2 or CDR 3 of the VKs (Fig. 3c, Group A, D and G), suggesting the light chain contributes to the RabMAb activity.

**Fig. 3a.**
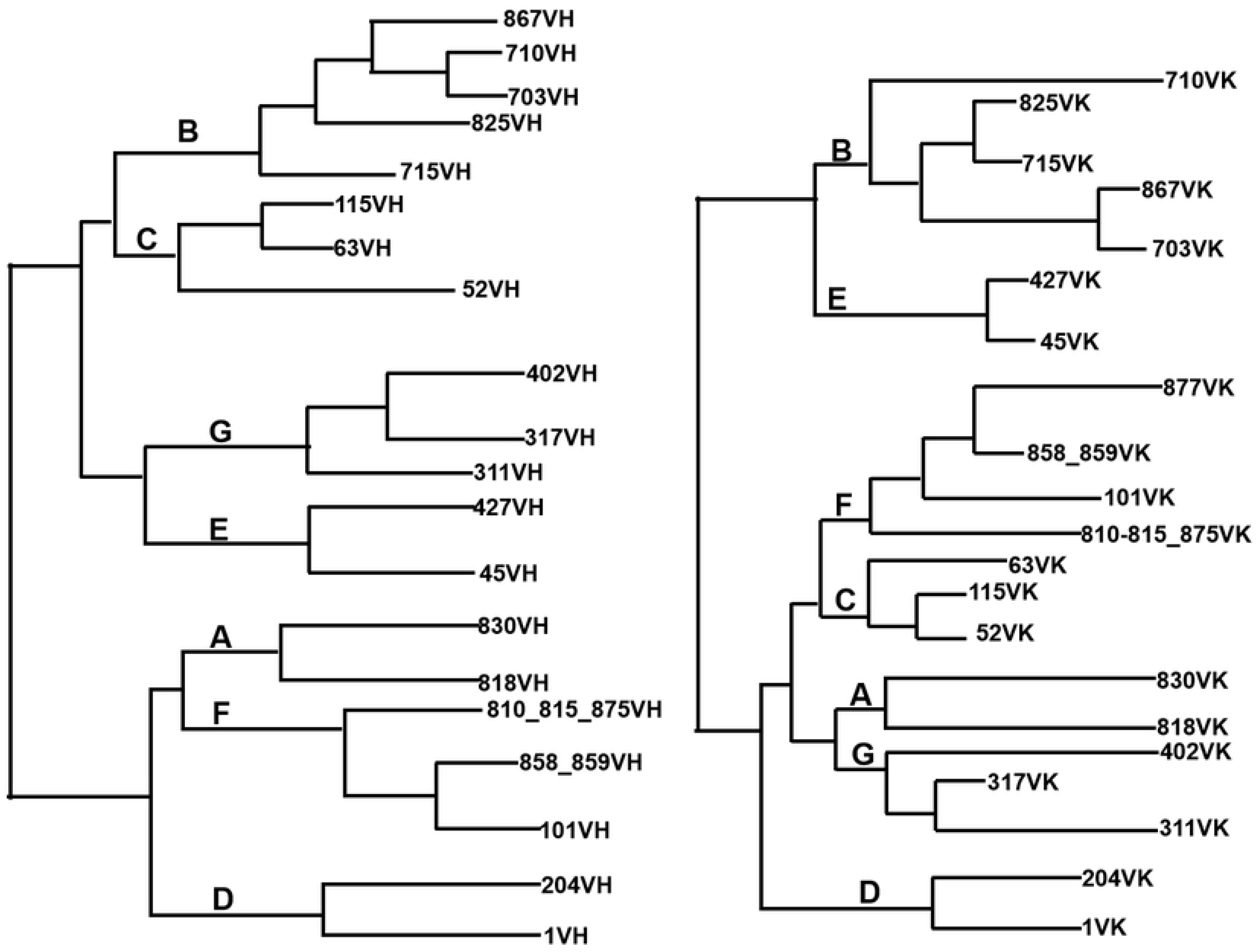
Lineage-related groups among the anti-TNFa RabMAbs

**Fig. 3b.**
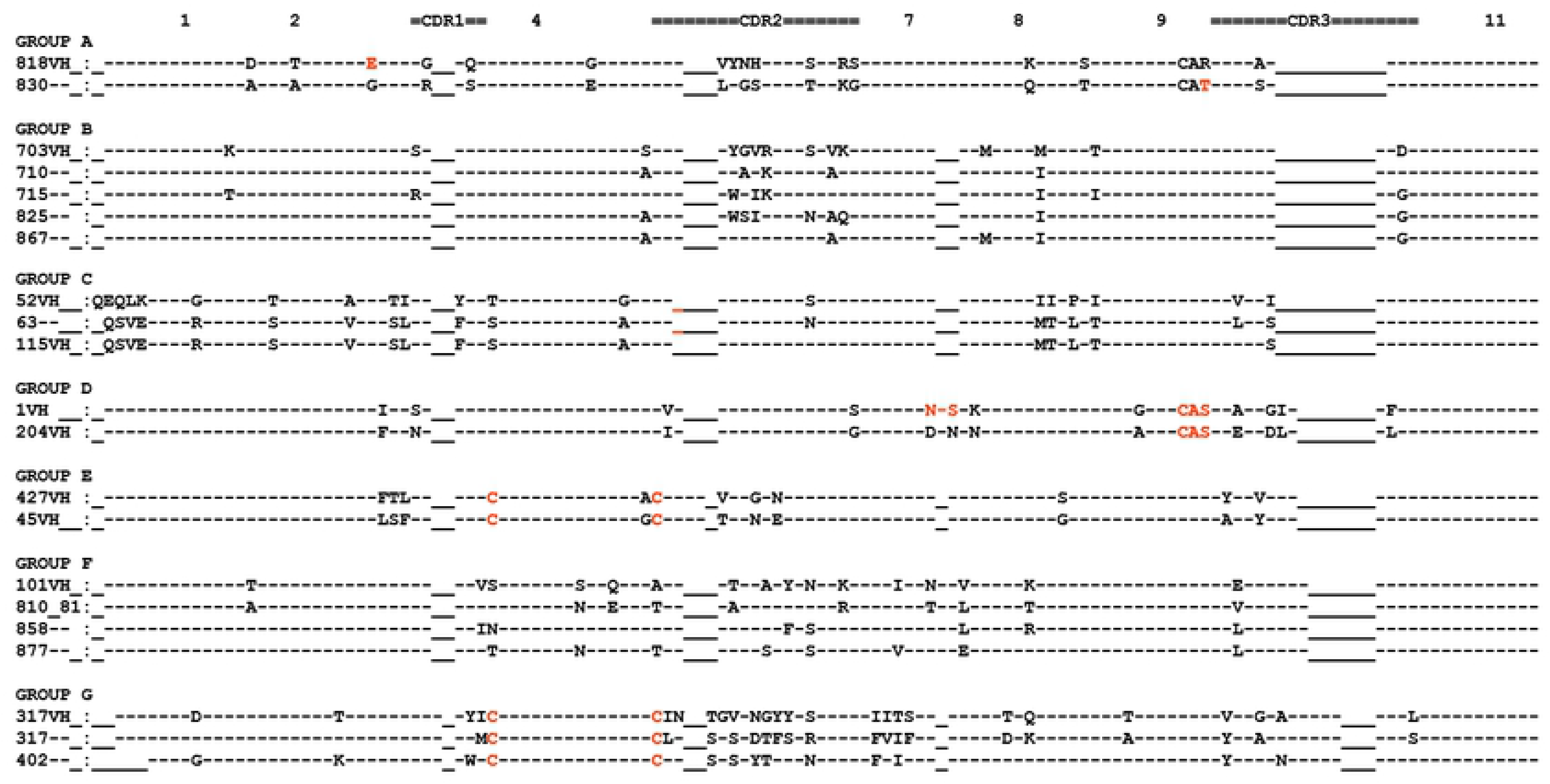
Lineage-related anti-TNFa RabMAb VH sequence alignment

**Fig. 3c.**
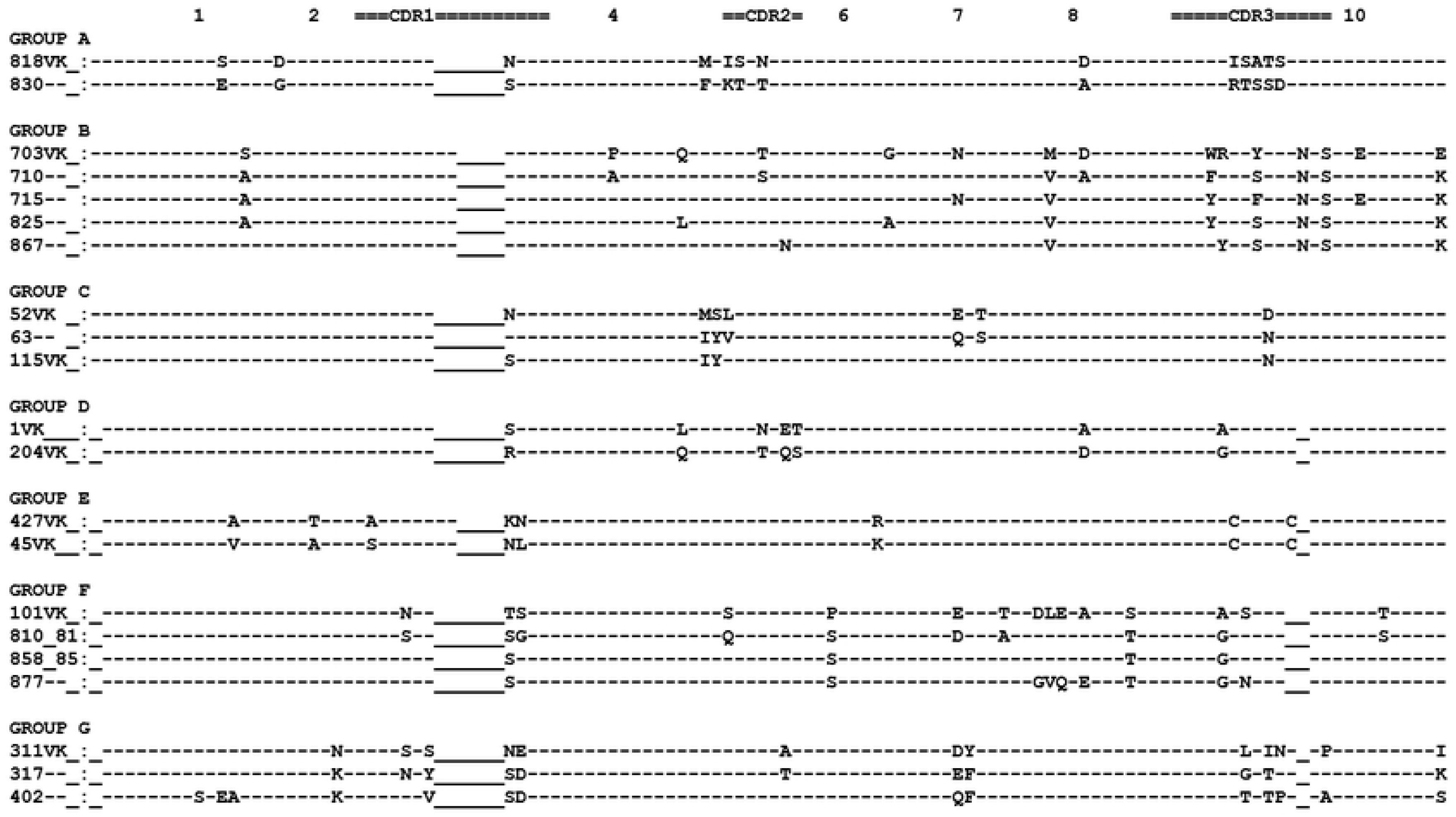
Lineage-related anti-TNFa RabMAb VK sequence alignment

Table 5 summarizes sequence and structure features of the anti-TNFa RabMAbs. Except for Clone 2 and 878, there is no corresponding human L3 canonical structure, and for some of the antibodies, no corresponding human H2 or L1 canonical structure either^11^. Some other uncommon sequence and structure features were also found, including extra Cys-Cys pairs in VH (CDR1 and CDR2) or VK (within L3) or both, lone Cys residue, potential glycosylaton sites, no CAR sequence in H92, 93 and 94, a deletion in H2 and no Gly residue at H26^12^.

**Table 5.**
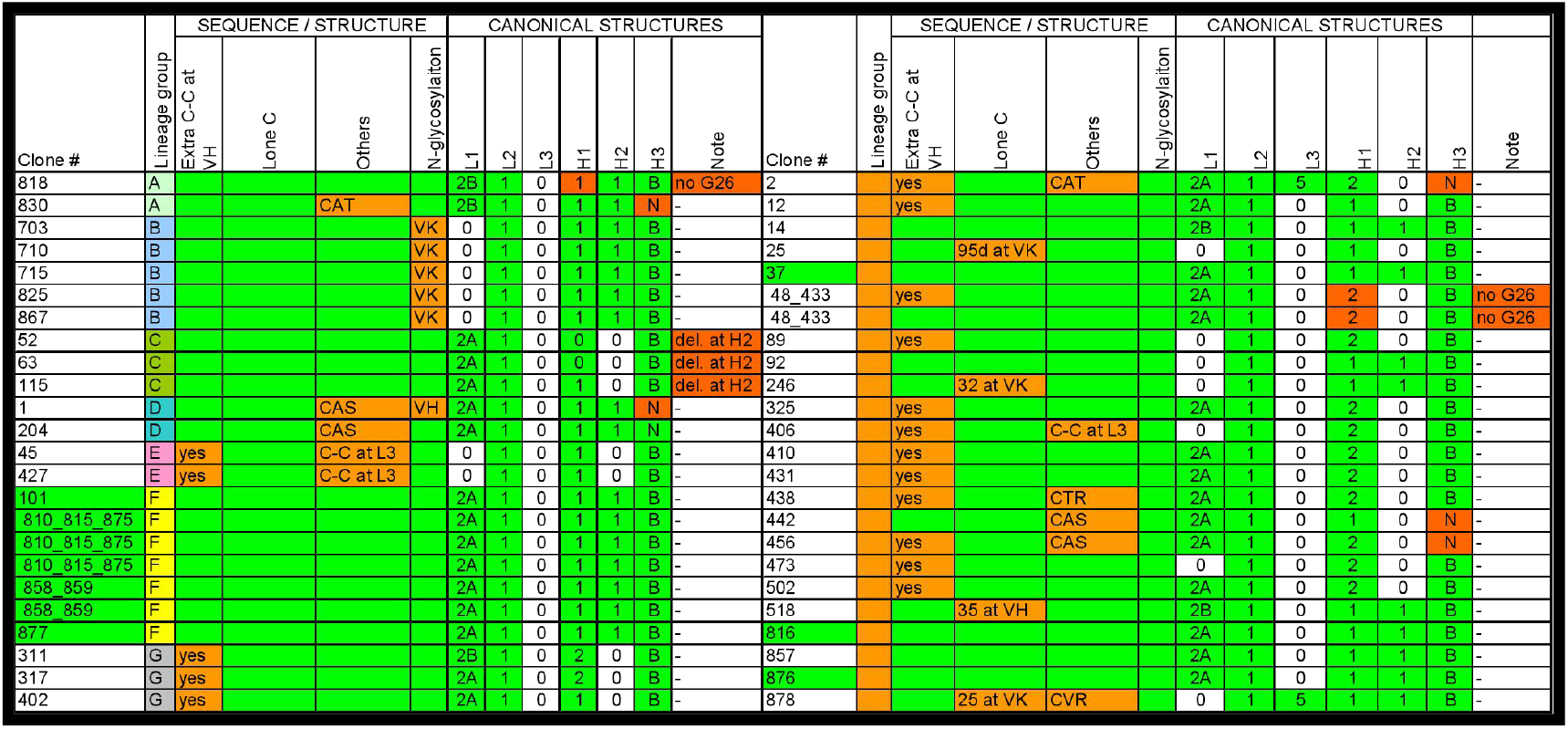
Summary of Sequence and Structure Analysis of Anti-TNFa RabMAbs

### Humanization of Anti-TNFa RabMAb Clone 858

In the lineage-related group F, all antibodies have a good neutralizing activity and antigen binding affinity (Table 4). In addition, the group F contains the most RabMAbs among the lineage-related groups, and does not have uncommon sequence and structure features that were found in other lineage-related groups or RabMAbs (Table 5). One RabMAb, Clone 858, in the group F was selected as a therapeutic lead candidate for humanization and further studies.

Clone 858 was humanized into a human IgG1 subclass with kappa light chain by using the MLG (mutational lineage-guided) method^5^. First, the heavy (VH) and light (VK) variable region sequences of Clone 858 were blasted against the human germnline VH and VK database. The closest human germline sequences, VH4-b/JH3 and VKIO2/JK4, were identified as the template for Clone 858 humanization (Fig. 4a and b). Second, antibody sequences within the Clone 858 lineage group (Fig. 3c and 3d, group F) were aligned to the template (Fig. 4a and b). Third, the rabbit residues in the frameworks potentially involved in CDR contacts or inter-chain contacts were identified based on the knowledge from human and mouse antibodies. These residues, marked by “#”, were not changed (1_, 2Q, 7F, 9L, 37V, 40A, 43S, 67S, 71R, 78V, 82M, 91F, 105P in VH and 1A, 2Y, 22K, 43P in VK) (Fig 4a and 4b). At CDR2 position 59 of VH, F is interchangeable to Y within the group, so it was mutated to the human residue Y at the position marked by * in Figure 4a. The same rule was applied to CDR1 position 28 of VK. N was changed to S at the position marked by * in Figure 4b. From the phylogenetic analysis, more residues in the frameworks were also considered not critical to structural activity of the antibody and humanized as marked by “$” (46Q – E of VH and 70E – D of VK). Residues at position 3D and 7T were substituted with human residue Q and S, respectively, as marked by “^^^” because they are away from the CDR loops and facing solvent (based on homology modeling). The parental Clone 858 antibody was 65% identical to the human germline frameworks (VH/VK). After humanization, the sequence identity is 90%.

**Fig.4.**
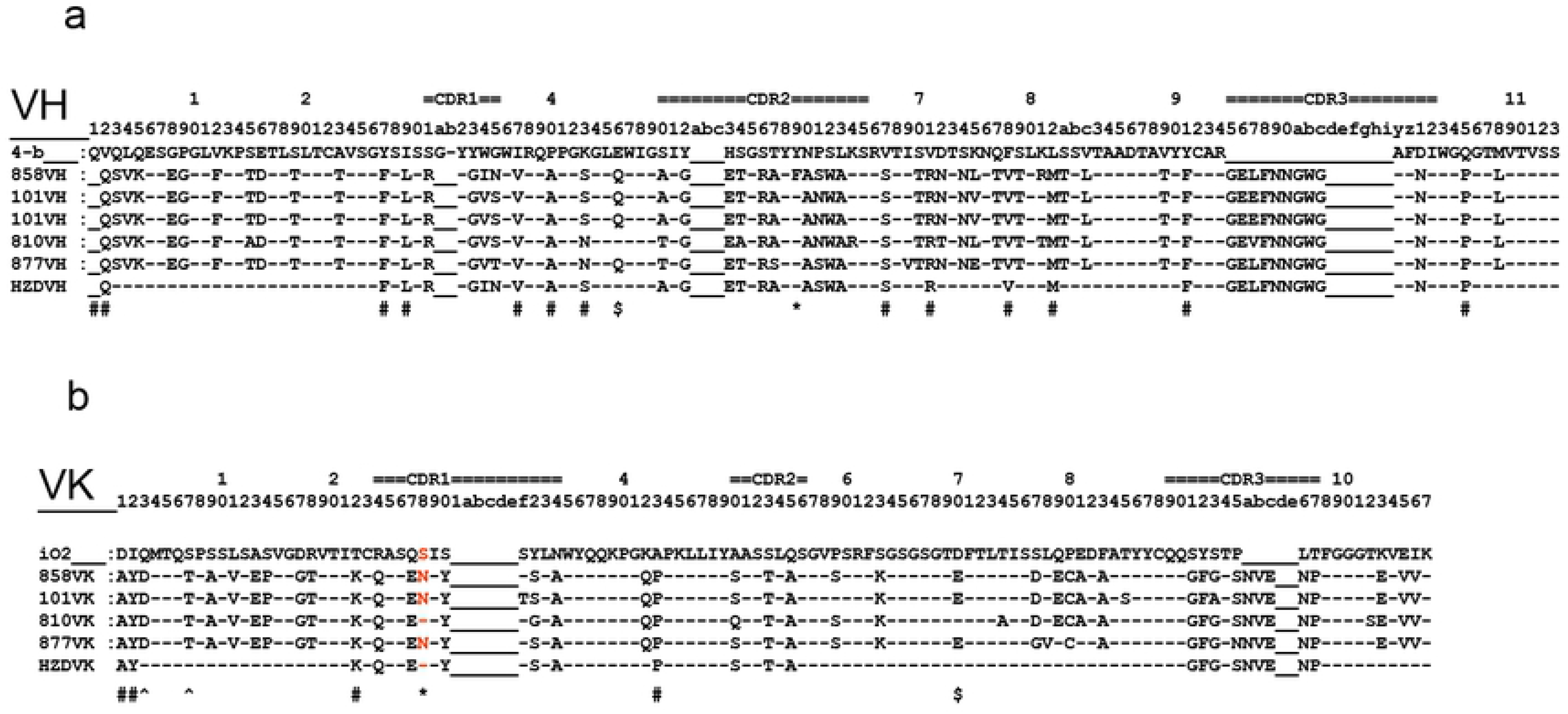
humanization of anti-TNFa Clone 858

### Comparison of Humanized Anti-TNFa RabMAb Clone 858 (HZD-RabMAb) to the Parental RabMAb and Marketed anti-TNFa drugs

To investigate whether HZD-RabMAb retains the properties of the parental antibody, we first compared N-glycosylation on the HZD and parental RabMAb and marketed anti-TNFa drugs. The HZD RabMAb, the parental, Humira and Remicade were treated with or without N-Glycosidase F (PNGase F), and analyzed by gel electrophoresis. Like Humira and Remicade, without PNCase treatnment, HZD-RabMAb only showed one heavy chain band (Fig. 5, at 50 KD). However, the parental RabMab showed two heavy chain bands, which is likely due to additional glycosylation site on the heavy chain CH1, predicted by sequence analysis, and partially gylcosyated product (Fig. 5, at 50 KD). Treatment with PNGase resulted in one deglycosylated heavy chain product for the HZD-RabMAb as well as for Humira, Remicade and the parental RabMAb (Fig.5, compare + PNGase to – PNGase). These results indicated HZD-RabMAb is the same as Humira and Remicade, but differentiated from the parental in glycosylation.

**Fig. 5.**
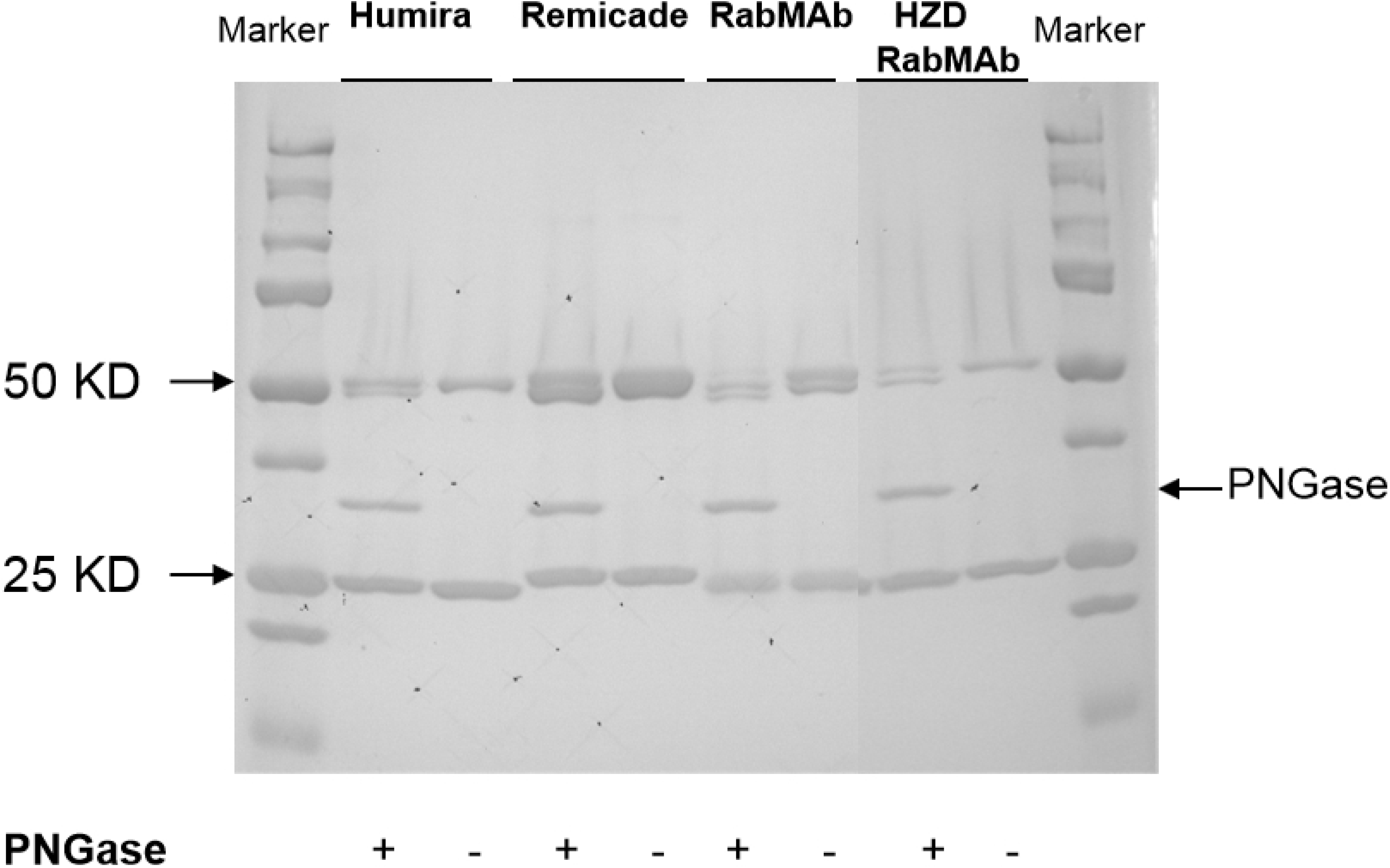
Deglycosylation of anti-TNFa antibodies. Anti-TNFa RabMAb Clone 858 and its humanized vesion, Humira and Remicade were treated with or without PNGase (N-glycosidase F), and then subjected to denatured, reducing gel electrophoresis. Proteins in the gel were visualized by Gelcode stain.

In addition to glycosylation, RabMab kappa light chain has a Cys-Cys pair between the framework 3 and the constant region (CK)^13^. Replacement of rabbit CK with human CK and substitution of the Cys with Pro in the framework 3 (Fig. 4b) for humanization could affect the light chain folding and make antibody structurally fragile. To address this issue, thermal stability of HZD-RAbMAb was examine and compared to Humira and Remicade. The thermal stability profile is shown in Fig. 6. HZD-RabMAb started its structure melting at about 74 °C as Humira and Remicade, indicating HZD-RabMAb is structurally as stable as the two marketed anti-TNFa drugs.

**Fig. 6.**
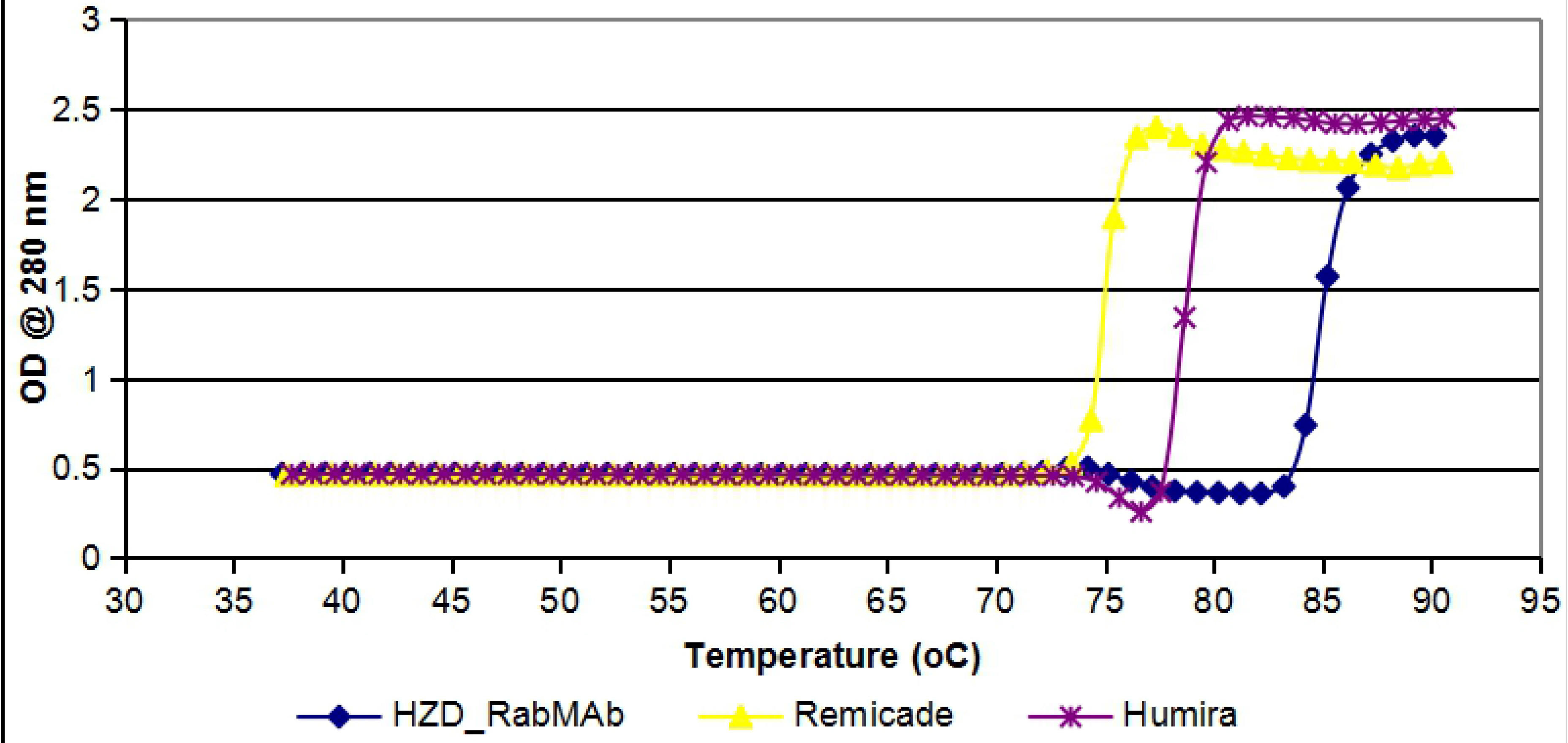
Thermal stability of anti-TNFa antibodies. Humanized anti-TNFa RabMAb Clone 858 (HZD-RabMAb), Humira and Remicade were subjected to thermal denaturation. Changes in absorbance at 280 nm was monitored over the temperature range of 37 to 90 ° C using a heating rate at 1 ° C/min.

Biological activity of HZD-RabMAb was tested for its TNFa neutralizing activity and binding activity. The relationship of L929 cell viability and antibody concentration is shown in Fig. 7. IC50 for HZD-RAbMAb is similar to that for the parental RabMab, but 1.5 times better than that (4.0 vs 9.0 ng/ml) for Remicade, indicating a high potency of HZD-RabMAb in neutralizing TNFa-induced cell toxicity. The antigen binding affinity measured by BIAcore is summarized in Table 6. Humanization of the RabMAb didn’t change its binding affinity to TNFa. Under the measurement conditions, HZD-RAbMAb showed picomolar affinity to the antigen.

**Fig. 7.**
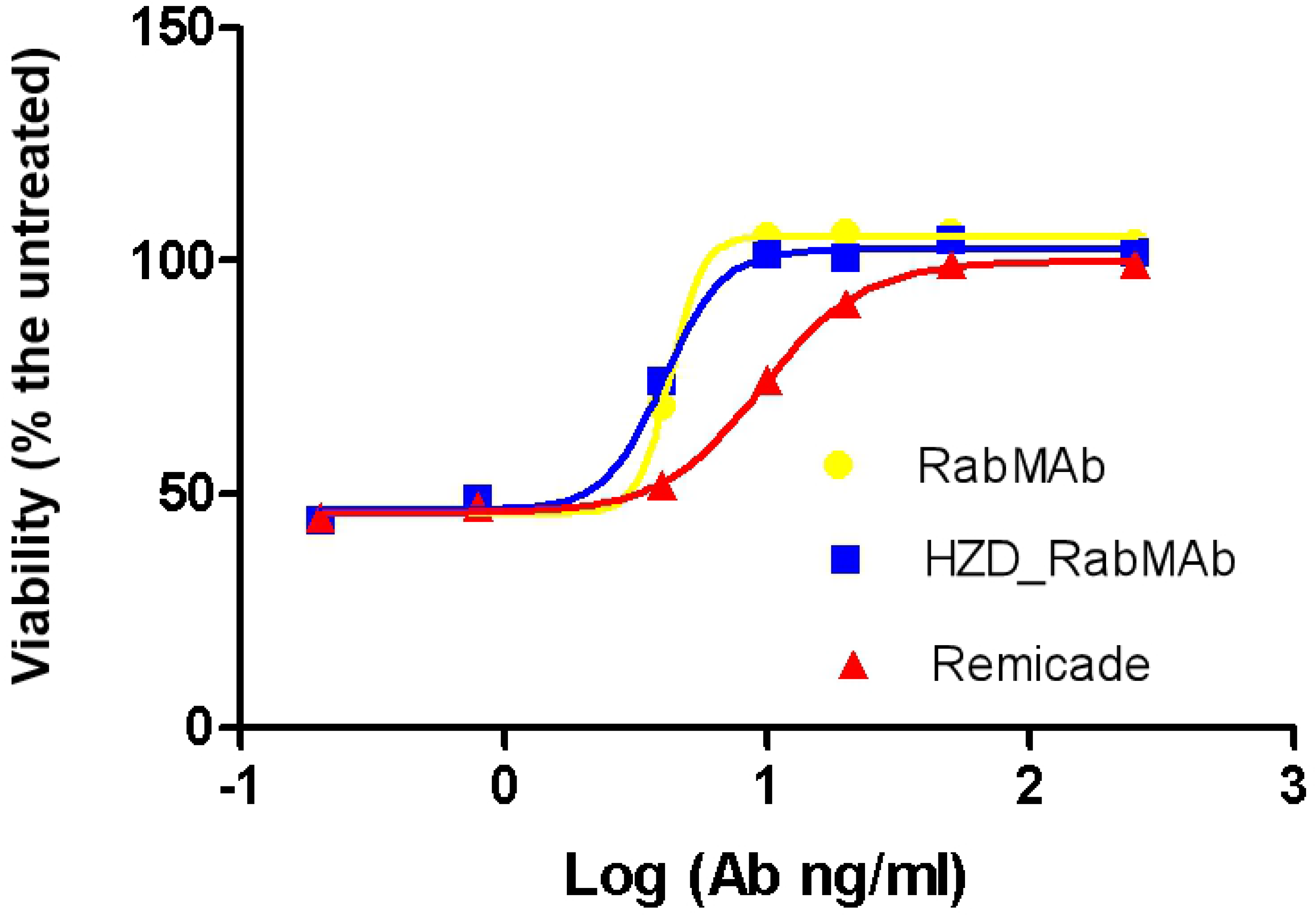
Neutralizing activity of anti-TNFa antibodies. L929 cells were treated with human TNFa and different concentrations of Anti-TNFa RabMAb Clone 858 (RabMAb), the humanized version (HZD-RabMAb), and Remicade, and then MTT assay was performed and absorbance at 570 nm was measured. The cells without or with TNFa treatment were run in parallel. Cell viability was calculated by normalizing the absorbance readings to no TNFa treatment. IC50 was determined by using GraphPad Prism 5 software

**Table 6.** Binding affinities of anti-TNFa HZD RabMAb and its parental RabMAb Clone 858*.

| Anitbody | Kd (pM) | Analyte conc. (nM) |
| --- | --- | --- |
| RabMab | 1.20±1.52 | 0.03-4 |
| HZD-RabMab | 1.49±0.46 | 0.05-3 |

\*Binding was performed in 10 mM HEPES, 150 mM NaCl, 0.05% P-20, pH 7.4 (HBS-P) at 35 μl/ml flow rate. 250 μl of analytes at various concentrations were injected and the dissociation lasts for 900 seconds. Data were fitted with 1:1 (Langmuir) binding model. The data were obtained from two measurements

### Pharmacokinetics Study

Humira, the RabMAb as well as the humanized RabMAb HZ3-M were injected to C57BL/6 mice intraperitoneally at 10mg/kg. The drug concentration in the animal blood was measured by ELISA at 5min, 30min, 1h, 4h 8h, 24h, 96h, 8d, 11d, 15d after the treatment. The result was shown in table 7 and figure 8. The elimination phase half-life for Humira, the humanized Rabmab HZ3-M and the Rabmab, was 105h, 167h, and 58.2h respectively, calculated using the single-compartment open model with first order absorption.

**Table 7.** Mice blood drug concentration at different time after i.p. injection.

| Time (h) | Serum Concentration (μg/ml) |  |  |
| --- | --- | --- | --- |
|  | HZ3M | RabMab | Humira |
| 0.0833 | -- | 1.54±0.93 | 2.76±0.35 |
| 0.5 | 14.89±5.00 | 3.62±4.56 | 14.89±11.31 |
| 1 | 26.84±11.38 | 21.71±7.73 | 16.50±7.89 |
| 4 | 93.90±33.53 | 95.66±7.69 | 50.06±21.27 |
| 8 | 58.54±20.64 | 67.22±14.57 | 48.10±18.54 |
| 24 | 81.30±23.46 | 49.48±7.33 | 59.15±7.15 |
| 96 | 68.10±12.58 | 23.20±3.72 | 49.58±4.95 |
| 192 | 39.40±15.44 | 13.04±2.72 | 26.08±7.50 |
| 264 | 27.93±12.17 | 1.40±1.45 | 13.67±9.11 |
| 360 | 22.73±9.46 | 1.48±0.88 | 9.11±5.70 |

**Figure 8.**
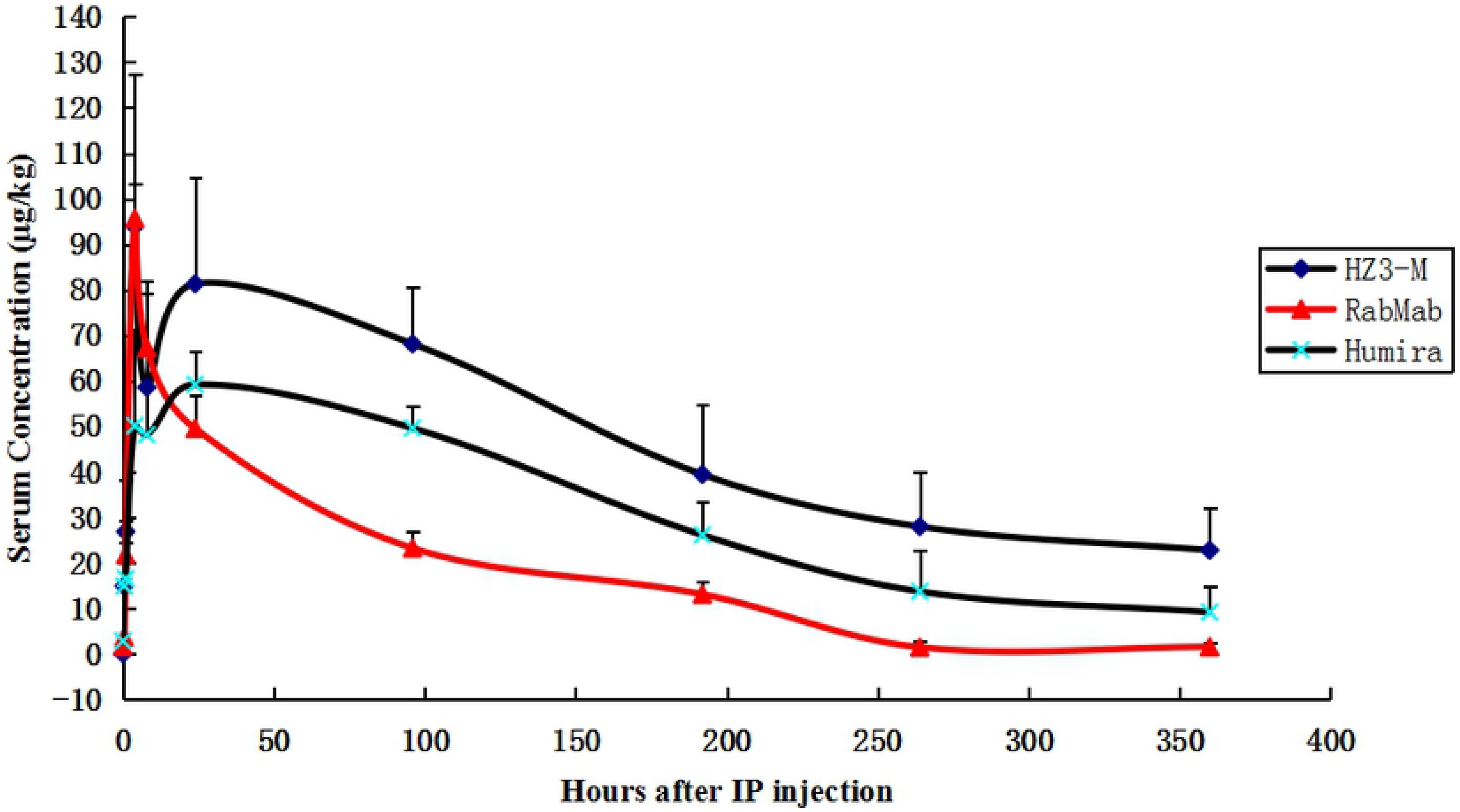
Mice blood drug concentration at different time after i.p. injection.

### Mice Efficacy Study

#### RA study

Normal mice (C57BL6) or human TNF-alpha transgenic mice (Taconic 1006) were with 1 mg/kg Humira (positive control), HZ3M at 0.33 mg/kg, 1 mg/kg, or 3 mg/kg, or solvent (negative control), as shown in table 1. All the treatment was given by i.p. injection, at 0.1 ml per 10 gram body weight once every three days for total of six weeks.

The animal arthritis symptoms were examined once a week after the first treatment and scored based on the limbs swollenness, toes curvature, toes separation, toe joints and ankle joints mal-shape, as described in the Materials and Methods. Humira was used as the positive control. The effect of HZ3M on normal and arthritis mice (Taconic 1006 mice) is shown in table 8 and figure 9.

**Table 8.**
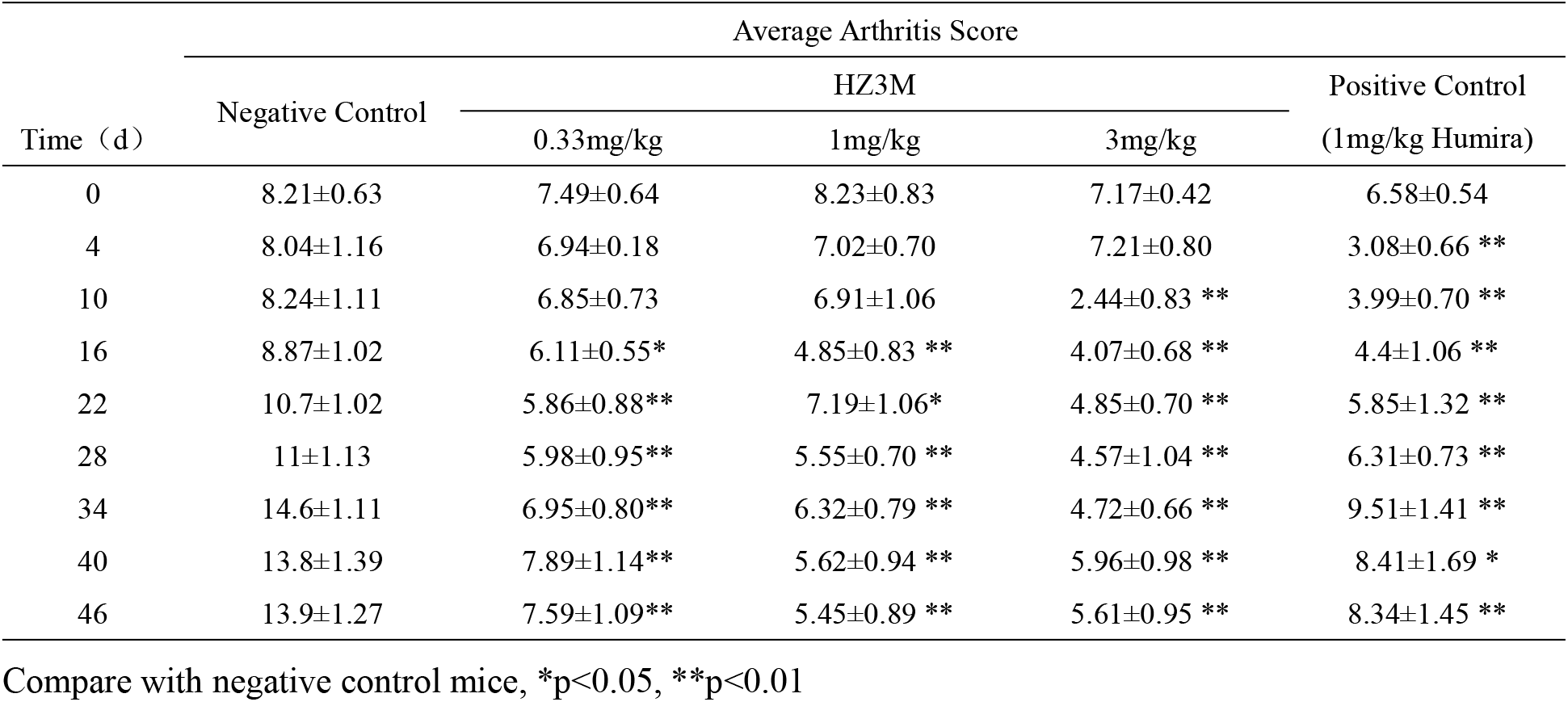
The effect of HZ3M on normal and arthritis mice (mean ±S.E.M)

**Figure 9.**
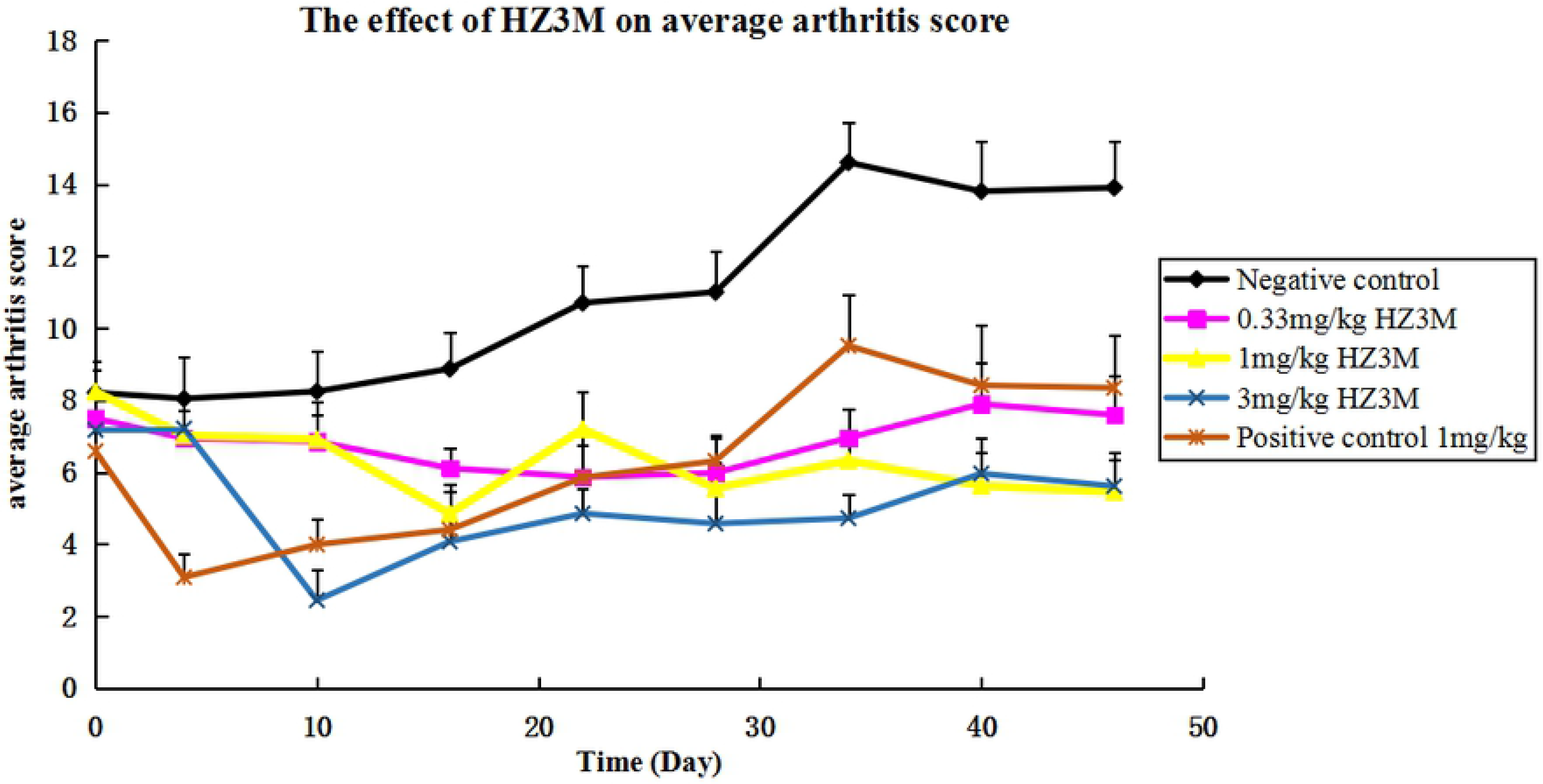
The effect of HZ3M on normal and arthritis mice

The arthritis symptoms were significantly reduced in Humira and HZ3M treated Taconic 1006 mice after two weeks of treatment.

#### Mice muscle strength study

The mice were subject to the muscle strength test once a week on the Rota-Rod Instrument during the treatment. The holding time was recorded, above 60 seconds were counted as 60 second.

During the treatment period, the Taconic 1006 mice suffer from the arthritis progression. The holding time was reducing. The arthritis symptoms in the Humira and HZ3M treated mice were significantly improved. Starting at ten days after the treatment, the Humira treated mice showed significantly longer holding time compared to the negative control mice (P< 0.05 at 10, 16, 34, 40; P<0.01 at day 22, 28, and 46). The HZ3M treated mice also showed significant longer holding time at all three doses, compared to the control group (for the high dose group, p<0.05 at day 10 and P<0.01 at day 16, 22, 28, 34, 40, and 46; for the medium dose group, P<0.05 at day 16 and P<0.01 at day 22, 28, 34, 40, and 46; for the low dose group, P<0.05 at day 40 and P<0.01 at day 22, 28, 34, 46). The final results are shown in table 9.

**Table 9.** The effect of Humira and HZ3M on muscle strength in the arthritis mice

| Time<br>(d) | Average Holding Time (sec.) |  |  |  |  |  |
| --- | --- | --- | --- | --- | --- | --- |
|  | Normal Mice | Taconic 1006 | Humira (1mg/kg) | HZ3M (0.33mg/kg) | HZ3M (1mg/kg) | HZ3M (3mg/kg) |
| 0 | 53.70±5.38 | 17.70±3.24 | 24.50±4.01 | 18.70±4.93 | 23.20±5.54 | 14.40±3.04 |
| 4 | 60.00±0.00 | 24.90±5.50 | 25.40±6.63 | 21.20±6.81 | 23.70±5.50 | 23.50±6.15 |
| 10 | 60.00±0.00 | 19.70±4.05 | 40.70±7.23* | 27.80±5.68 | 26.50±6.37 | 38.40±5.50* |
| 16 | 58.90±1.10 | 19.50±3.16 | 39.20±6.91* | 25.40±3.77 | 40.20±6.62* | 39.00±5.54** |
| 22 | 60.00±0.00 | 16.60±3.72 | 45.80±7.25** | 50.70±4.68** | 54.30±4.54** | 56.50±3.08** |
| 28 | 60.00±0.00 | 14.70±3.04 | 38.90±7.81** | 48.10±5.87** | 43.60±6.12** | 51.00±5.01** |
| 34 | 59.07±0.30 | 16.00±3.64 | 35.30±7.53* | 41.60±6.54** | 44.80±4.98** | 50.50±4.02** |
| 40 | 55.80±2.82 | 15.30±4.71 | 34.50±7.68* | 31.00±5.78* | 42.90±6.97** | 55.50±2.70** |
| 46 | 58.20±1.28 | 15.50±5.76 | 40.70±5.94** | 41.50±4.96** | 44.60±6.55** | 46.90±4.96** |

Compare with negative control mice, \*p<0.05, \*\*p<0.01

**Figure 10.**
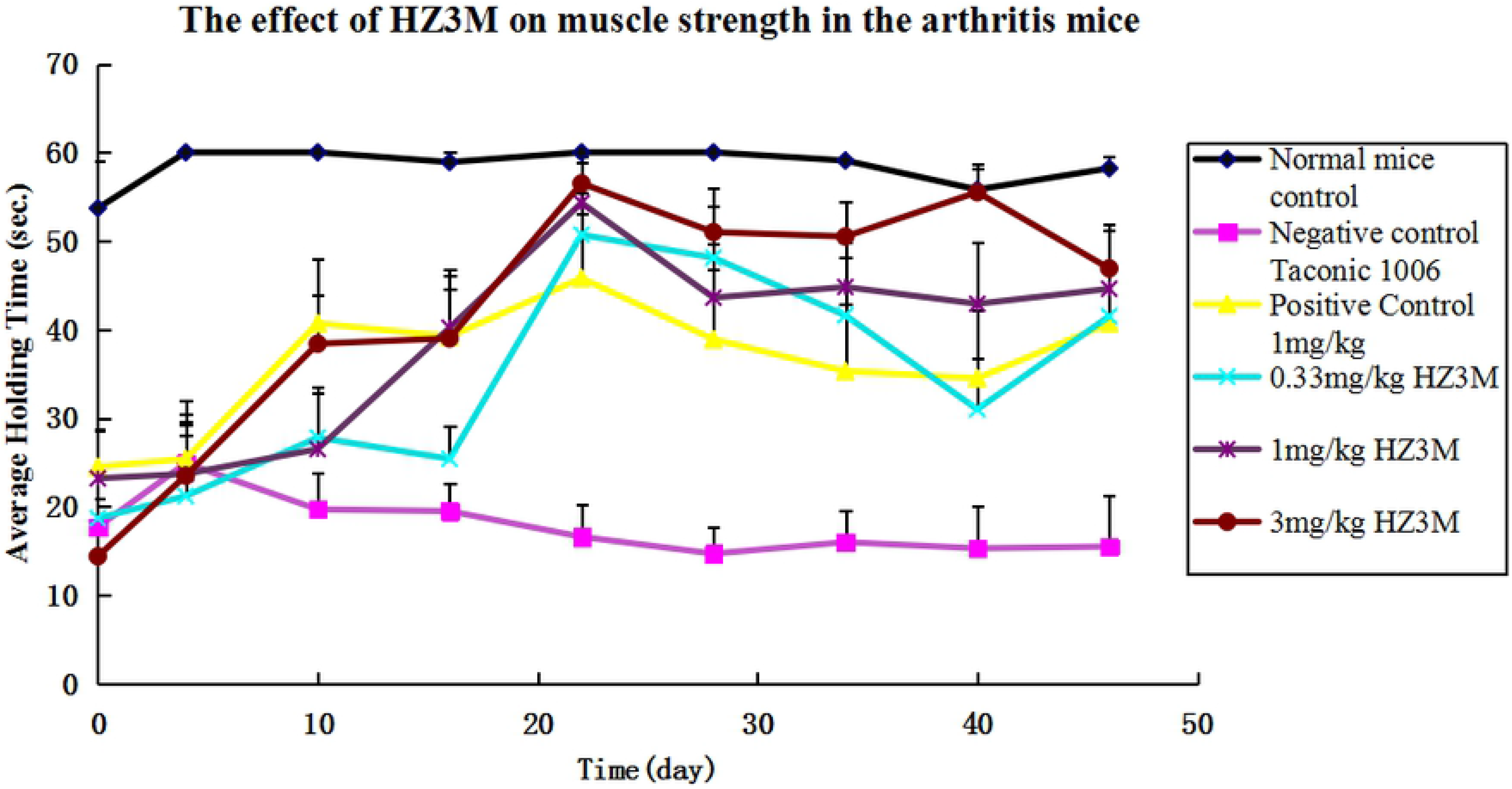
The effect of Humira and HZ3M on muscle strength in the arthritis mice

#### Joint histology analysis

the results of histology scores of treated animals were shown in table 10.

**Table 10.**
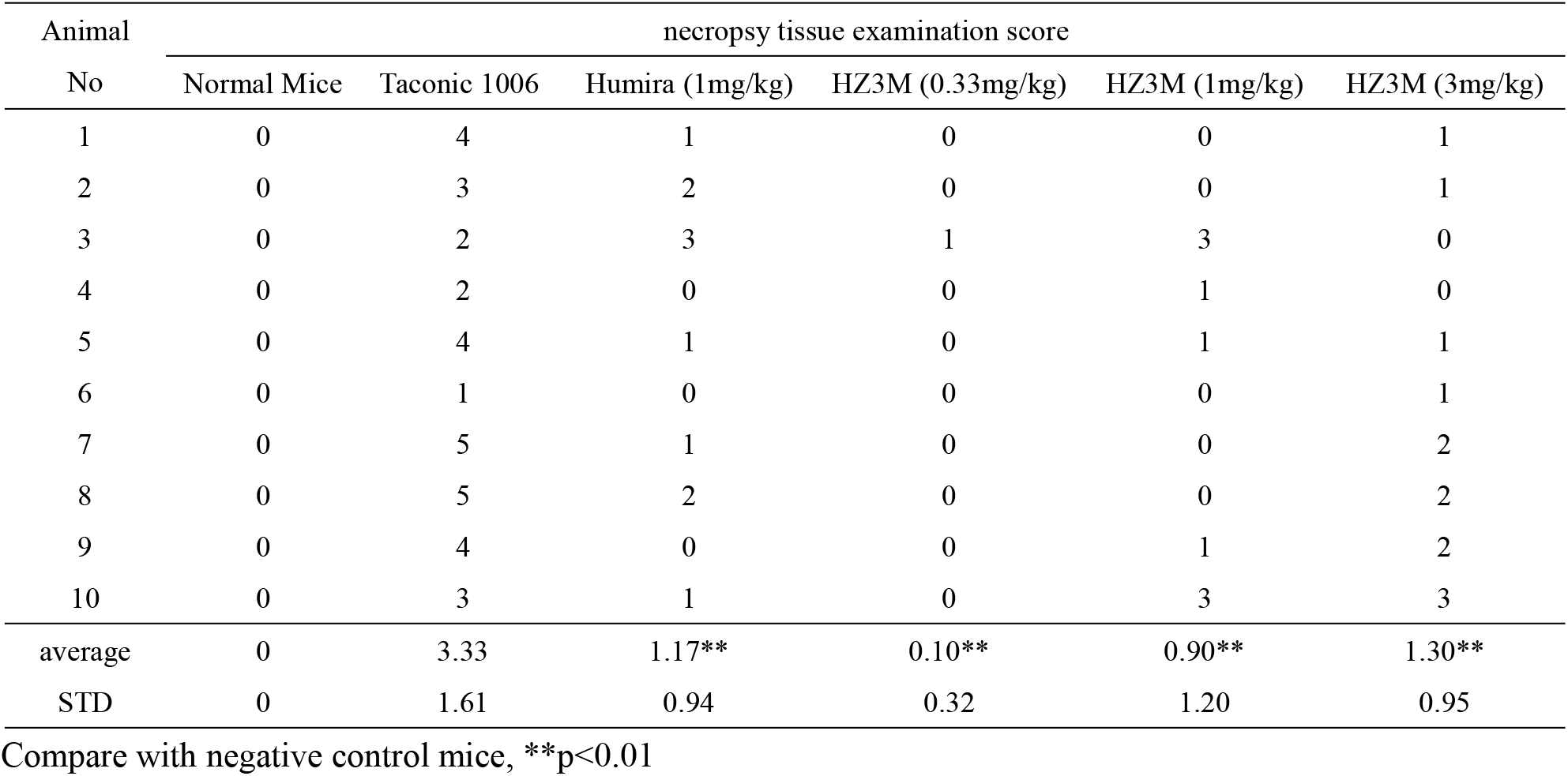
The effect of Humira and HZ3M on microscopically necropsy tissue examinein the arthritis mice

### Mice survival study

The Normal mice did not survive from the deadly dose of hTNFα with effect enhancer D-galactosamine. With the protection of anti-TNFα monoclonal antibodies, mice death rate is decreased. The anti-TNFα monoclonal antibodies do not just bind to TNFα but also block the interaction between TNFα and its receptors, so the bioactivities of TNFα can be neutralized. In the HZ3M group, the death rates are decreased with those increasing doses. A clear dose-dependent relationship between mice survival and HZ3M administration dose is seen in the table 11.

**Table 11.** The effect of Humira and HZ3M on mice survival study

| Group | Survival No./Total | Survival rate (%) |
| --- | --- | --- |
| Normal Mice | 10/10 | 100 |
| Negative Control | 1/10 | 10 |
| Positive Control Humira 0.10µg/g | 8/10 | 80 |
| HZ3M 0.07µg/g | 5/10 | 50 |
| HZ3M 0.10µg/g | 9/10 | 90 |
| HZ3M 0.13µg/g | 10/10 | 100 |

### Fever study

hTNFα is able to increase the temperature of rabbit with D-Gln. The anti-TNFα antibodies have neutralized the bioactivities of TNFα after administrated to rabbits. And like those in mice survival study, dose-dependent behaviour is found in the rabbit fever study as well (table 12 and figure 11).

**Table 12.** The effect of Humira and HZ3M on rabbit fever study (Mean±STD)

| $^{\circ}\text{C}$ \diagup min | 0 | 30 | 60 | 90 | 120 | 150 | 180 | 210 | 240 |
| --- | --- | --- | --- | --- | --- | --- | --- | --- | --- |
| Blank control group | 38.80±<br>0.29 | 39.30±<br>0.27 | 40.12±<br>0.4 | 40.02±<br>0.38 | 39.62±<br>0.27 | 39.32±<br>±0.48 | 39.05±<br>±0.59 | 38.98±<br>±0.5 | 38.76±<br>±0.46 |
| Negative control group | 38.78±<br>0.16 | 38.65±<br>0.12 | 38.65±<br>0.18 | 38.68±<br>0.17 | 38.67±<br>0.13 | 38.68±<br>±0.15 | 38.61±<br>±0.29 | 38.59±<br>±0.25 | 38.61±<br>±0.26 |
| Positive control group<br>(Humira 50µg/kg) | 38.80±<br>0.33 | 38.83±<br>0.38* | 38.83±<br>0.42** | 38.81±<br>0.49** | 38.87±<br>0.51** | 38.86±<br>±0.50 | 38.85±<br>±0.56 | 38.82±<br>±0.60 | 38.86±<br>±0.66 |
| HZ3M 25µg/kg | 38.77±<br>0.56 | 38.74±<br>0.53* | 39.55±<br>0.31** | 39.58±<br>0.26** | 39.11±<br>0.49** | 38.98±<br>±0.55 | 38.95±<br>±0.53 | 38.94±<br>±0.56 | 38.86±<br>±0.57 |
| HZ3M 50µg/kg | 39.00±<br>0.41 | 39.14±<br>0.42 | 39.18±<br>0.36** | 39.13±<br>0.36** | 39.16±<br>0.38* | 39.21±<br>±0.33 | 39.25±<br>±0.30 | 39.19±<br>±0.30 | 39.11±<br>±0.24 |
| HZ3M 100µg/kg | 38.86±<br>0.32 | 38.86±<br>0.29 | 38.85±<br>0.30* | 38.88±<br>0.35* | 38.99±<br>0.33* | 38.99±<br>±0.35 | 38.98±<br>±0.34 | 38.91±<br>±0.33 | 38.84±<br>±0.29 |

Compare with negative control mice, \*p<0.05, \*\*p<0.01

**Figure 11.**
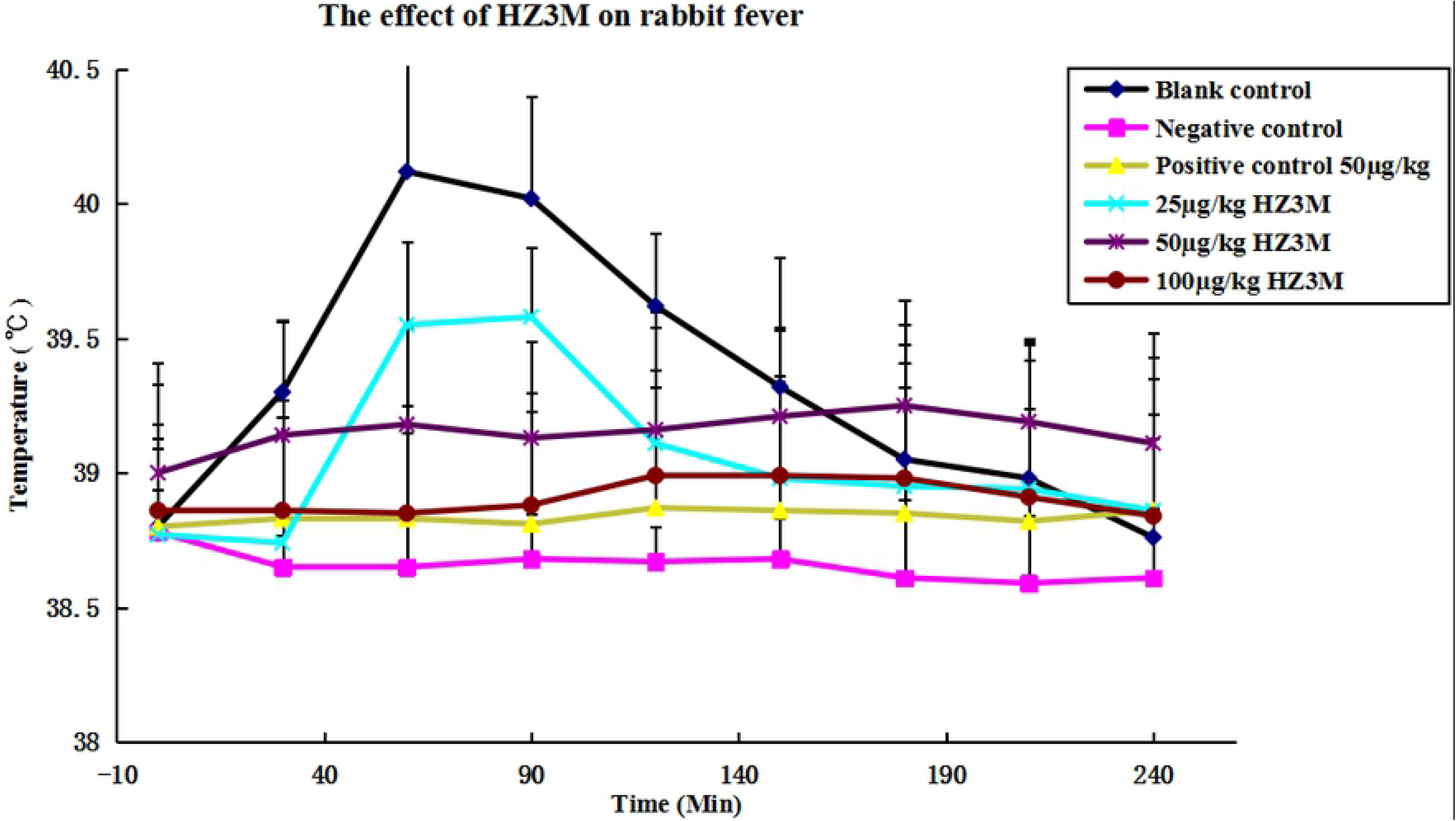
the effect of HZ3M on rabbit fever

## DISCUSSION AND CONCLUSIONS

In this study, we sought to develop a new anti-TNFa monoclonal drug, with high binding and neutralizing activity based on RabMAbs platform. Using MLR strategy, the humanization version of anti-TNFα RabMAb HZD-RabMAb (also named as HZ3M) is produced. The humanized RabMAb inherited most the activities of binding and neutralizing and the thermal stability. In vivo PK study, the behaviour of HZ3M is similar with its RabMAb version and two marketed anti-TNFa drugs. As expected with any formulation of HZ3M, it will be more safety to develop a humanied antibody, multiple experiments from our laboratories have primarily observed the effects of HZ3M on RA.In vivo efficay studies show HZ3M is able to bind and neutralize hTNFα in transgenic and normal mice as well as normal rabbits. Clearly dose-dependent response can be determined, compared to marketed anti-TNFa drug Humira, the efficacy of HZ3M is seems to be better, HZ3M was easy to see the advantages of the new antibody, the hafe-life was 167 hours, longer than drugs on the market Humira, which was 105 hours as a contral drug in our Pharmacokinetics study. The survival rate reasch showed a clear dose-dependent relationship between the mice survival rate and HZ3Madministration dose, in 0.1μg/g dose, it seems better survival rate than Humira, further more, in 0.13 μg/g dose, the survival rate was up to 100%. The survival rate further supports the administration of HZ3M to patients RA.These results encourage the further development of HZ3M derived from RabMAb to preclinical safety study.

